# Non-essential ribosomal proteins in bacteria and archaea identified using COGs

**DOI:** 10.1101/2021.01.31.429008

**Authors:** Michael Y. Galperin, Yuri I. Wolf, Sofya K. Garushyants, Roberto Vera Alvarez, Eugene V. Koonin

## Abstract

Ribosomal proteins (RPs) are highly conserved across the bacterial and archaeal domains. Although many RPs are essential for survival, genome analysis demonstrates the absence of some RP genes in many bacterial and archaeal genomes. Furthermore, global transposon mutagenesis and/or targeted deletion showed that elimination of some RP genes had only a moderate effect on the bacterial growth rate. Here, we systematically analyze the evolutionary conservation of RPs in prokaryotes by compiling the list of the ribosomal genes that are missing from the one or more genomes in the recently updated version of the Clusters of Orthologous Genes (COG) database. Some of these absences occurred because the respective genes carried frameshifts, presumably, resulting from sequencing errors, while others were overlooked and not translated during genome annotation. Apart from these annotation errors, we identified multiple genuine losses of RP genes in a variety of bacteria and archaea. Some of these losses are clade-specific, whereas others occur in symbionts and parasites with dramatically reduced genomes. The lists of computationally and experimentally defined non-essential ribosomal genes show a substantial overlap, revealing a common trend in prokaryote ribosome evolution that could be linked to the architecture and assembly of the ribosomes. Thus, RPs that are located at the surface of the ribosome and/or are incorporated at a late stage of ribosome assembly are more likely to be non-essential and to be lost during microbial evolution, particularly, in the course of genome compaction.

**IMPORTANCE:** In many prokaryote genomes, one or more ribosomal protein (RP) genes are missing. Analysis of 1,309 prokaryote genomes included in the COG database shows that only about half of the RPs are universally conserved in bacteria and archaea. In contrast, up to 21 other RPs are missing in some genomes, primarily, tiny (<1 Mb) genomes of host-associated bacteria and archaea. Ten universal and nine archaea-specific ribosomal proteins show clear patterns of lineage-specific gene loss. Most of the RPs that are frequently lost from bacterial genomes are located on the ribosome periphery and are non-essential in *Escherichia coli* and *Bacillus subtilis*. These results reveal general trends and common constraints in the architecture and evolution of ribosomes in prokaryotes.

## INTRODUCTION

Ribosomes are macromolecular cell factories that consist of rRNAs and ribosomal proteins (RPs) and are responsible for the translation of all mRNAs. Bacterial ribosomes that have been thoroughly characterized in model organisms, such as *Escherichia coli* and *Bacillus subtilis*, typically contain 54 core RPs, 33 in the large subunit and 21 in the small subunit (1–5). Archaeal ribosomes include up to 66 proteins, of which 33 are universal, i.e. shared with bacteria and eukaryotes (18 in the large ribosomal subunit and 15 in the small subunit), and 33 proteins are shared only with eukaryotes. The list of core RPs in several model organisms is provided in Supplementary Table S1.

Several lines of evidence indicate that some RPs can be non-essential, at least, in some organisms and under certain conditions. First, experiments on genome-wide mutagenesis have resulted in the generation of mutants with a deletion or transposon insertion in a variety of RP genes. Such mutants were viable, albeit grew slower than the wild type (6, 7). Such experiments have been performed in a wide variety of bacteria, albeit, so far, not in archaea. The global mutagenesis approach has some potential caveats, such as conditional lethality (mutations in two genes may be tolerated individually but not together) and functional compensation by paralogs. For example, *E. coli* encodes paralogs of L31 and L36, whose expression can suppress the lack of these RPs (8). Similarly, *B. subtilis* encodes two paralogs of L31, L33, and S14 each that could partly compensate for the loss of the respective RP function (9, 10). In addition, the absence of certain RP genes can be compensated by changes in the intracellular milieu, such as, for example, high level of Mg^2+^ ions (11, 12). Gene essentiality data derived from genome-wide mutagenesis studies are well represented in the literature and are also available in such online databases as the Database of Essential Genes [DEG, (13)] and the Online Gene Essentiality database [OGEE, (14)]. In addition to the global mutagenesis studies, data on gene essentiality have been obtained by monitoring the effects of suppressing gene expression, e.g. with antisense RNA (15–17).

Another general approach to the prediction of (non)essential RPs is by using comparative genomics (1–5). The absence of a particular gene in a complete microbial genome (or, better yet, in several related genomes) strongly suggests that this gene is non-essential, at least for growth on a rich medium. This approach also has several caveats, such as the problems with genome completeness and sequencing quality, as well as the presence of paralogs or other forms of functional compensation. However, it is cheap, high-throughput, and readily applies to hard-to-grow and even non-cultured bacteria and archaea. Genome comparisons have proved particularly fruitful for the analysis of the highly reduced genomes of intracellular parasites, insect cell symbionts, and for the near-minimal genomes of axenically-growing mollicutes (18–25). Collectively, these studies suggest that the number of truly essential RP genes could be as small as 33 (21).

The universal presence of most RPs in organisms from all three domains of life makes them a key component of the small set of highly conserved genes that can be used for the construction of deep-rooted phylogenetic trees and the global Tree of Life (4, 26, 27). Therefore, understanding the evolution of RPs and differentiating universal, essential RPs from dispensable ones that are occasionally lost during evolution is an important task in evolutionary biology.

Here, we report the patterns of presence and absence of RP genes in the current release of the Clusters of Orthologous Genes (COGs) database (28). The COG database is a particularly convenient tool for the analysis of gene gain and loss because it includes a limited number of high-quality complete microbial genomes and features COG-specific patterns of presence and absence of evolutionarily conserved genes in the respective organisms (29–31). In addition, the COG construction algorithm (32, 33) provides for the detection of even highly diverged orthologous proteins that are not necessarily recognized as orthologs by other tools (30, 31, 34, 35). Phyletic patterns of COGs have been previously used to reconstruct the ancestral states and evolution of various functional systems including the minimal and ancestral sets of RPs (2, 36). Owing to these features, the COG database allows for straightforward identification of the genomes that do not encode the given RP.

The current version of the COG database (28) features a selection of COGs grouped by metabolic pathways and functional complexes, including the RPs of the 50S and 30S ribosome subunits as well as a group of archaea-specific RPs. Examination of the phyletic patterns of the COGs for all three groups allowed us to (i) compile the list of about 500 RP genes missing in some bacterial and/or archaeal genomes (some actually lost and some missing because of sequencing problems); (ii) identify more than 50 RP genes that have been overlooked in the course of genome annotation, and (iii) establish the patterns of RP gene loss during bacterial and archaeal evolution, and iv) correlate the experimentally-derived and computationally-generated sets of the likely non-essential RP genes.

## RESULTS

### Delineation of the ribosomal protein set

The conserved ribosomal protein (RP) set, extracted from the current release of the COG database (28), consisted of 54 core RPs: 33 from the 50S subunit (L1-L7/L12, L9-L11, L13-L25, and L27-L36) and 21 (S1-S21) from the 30S subunit (1–5). Several additional proteins, such as S22 (RpsV, Sra) and S31e (Thx), which are associated with ribosomes in some bacteria (37, 38), are not covered in the COG database and have not been included in the analyzed set. The archaeal RP gene set included 66 genes, of which 33 are shared with bacteria and eukaryotes (18 in the large ribosomal subunit and 15 in the small subunit), and 33 RPs that are shared only with eukaryotes. The list of of core RPs from model organisms such as *Escherichia coli* K-12, *Bacillus subtilis* strain 168, *Mycoplasma pneumoniae* M129, *Aeropyrum pernix* K1, and *Haloarcula marismortui* ATCC 43049, that were analyzed here is presented in Supplementary Table S1. This table shows that the archaea-specific RP set is quite variable: *A. pernix* encodes seven RPs that are missing in *H. marismortui*.

### Frameshifted and unannotated ribosomal protein genes

Before analyzing the patterns of RP loss across the diversity of bacteria and archaea, it was necessary to identify and eliminate artifacts that could result from sequencing or annotation errors. To ensure the quality of the genome collection, members of the International Nucleotide Sequence Database Collaboration, the DNA Database of Japan, EBI’s European Nucleotide Archive, and NCBI’s GenBank, routinely check new genome submissions for the presence of certain RPs (39). Nevertheless, due to the sheer number of sequenced genomes, errors occasionally crop up, which becomes evident when the same organisms repeatedly show up as missing certain RPs despite having relatively large genomes and in the absence of similar problems in related organisms. Another tell-tale sign of sequencing problems is the presence of frameshifted genes that are present in a full-length form in other members of the same lineage (Table S2). There are good reasons to suspect that many if not most of these frameshifts represent sequencing errors, rather than genuine mutations or cases of programmed translational frameshifting that is not known to be a common mechanism of RP translation (40). For example, the 6.09-Mb genome of the betaproteobacterium *Mitsuaria* sp. str. 7 misses the genes for L13, L21, L25, L27, and S9 proteins, which is unique among the genomes of this size. Likewise, the 3.97-Mb genome of the alphaproteobacterium “*Candidatus (Ca.)*. Filomicrobium marinum Y” lacks the L1, L7/L12, L10, L11, S7, and S12 genes (Table S2) and is the only genome where the genes for the latter two proteins are missing.

Another widespread cause of missing RPs is the automated genome annotation, which sometimes fails to recognize genuine protein-coding genes, particularly short ones. As a result, these overlooked ORFs are not included in the respective protein sets. Sequencing and annotation problems often show up in the same genomes, making their quality suspect and putting into question the apparent absence of certain RPs. As an example, in the 4.28-Mb GenBank entry for the halophilic gammaproteobacterium *Salinicola tamaricis* F01, *rplF* (encoding the L6 protein), *rplI* (L9), *rplL* (L7/L12), *rplY* (L25), and *rpsD* (S4) genes are frameshifted, the *rpsB* (S2) gene is absent, and two full-length genes, *rplD* (L4) and *rpsR* (S18), are present but have been overlooked in the course of genome annotation. Similarly, the current GenBank entry for *Sulfobacillus acidophilus* str. TPY, a member of the *Clostridia*, lacks the genes for RPs L27, L28, L32, L33, L36, and S14, which are encoded in the genome but have been overlooked in course of annotation. These genes have been easily found by the TBLASTn search (41) using the respective RPs from closely related clostridial genomes as queries (see Table S3 for details). Two more organisms, *Pelotomaculum thermopropionicum* and “*Ca.* Methylo-mirabilis oxyfera”, had five overlooked ribosomal genes each (Table S3). Two or more unannotated RP genes were found in seven more bacterial genomes.

The tiny (606-kb) genome of the nanoarchaeon “*Ca.* Nanopusillus acidilobi” presented a different problem. The current protein set of “*Ca.* Nanopusillus acidilobi” in GenBank misses 14 RPs that are found in almost all other archaeal genomes. However, a detailed examination of this genome showed that only four RP genes were truly missing (Table 1), the gene encoding S14 protein was frameshifted (Table S2), and the gene for L37e has been overlooked and could be found by TBLASTn (Table S3). Full-length ORFs coding for eight other RPs: L6p/L9e (genomic locus tag Nps_02895), L15e (Nps_01385), L16/L10ae (Nps_03305), L22 (Nps_03365), L24 (Nps_02910), L35ae (Nps_03205), S6e (Nps_01880), and S15p/S13e (Nps_01520), were correctly identified at the annotation stage and described in the respective publication (42). However, for some unknown reason, these genes were assumed to be disrupted and were erroneously marked as pseudogenes in the GenBank entry for “*Ca.* Nanopusillus acidilobi” (see Table S3 for details). As a result, the RPs encoded by these genes, which are all longer than 110 amino acids, never made it into the protein database. The same problem on a lesser scale was observed for the other nanoarchaeon in the current COG collection, “Nanohaloarchaea archaeon SG9”, where the genes for L24e, L40e, and S28e proteins were overlooked, whereas genes encoding L18 and S2 were marked as pseudogenes and left untranslated (Table S3). Correcting such annotation problems is important for assessing the essentiality of RPs in biologically interesting but poorly studied groups of microorganisms.

**Table 1.**
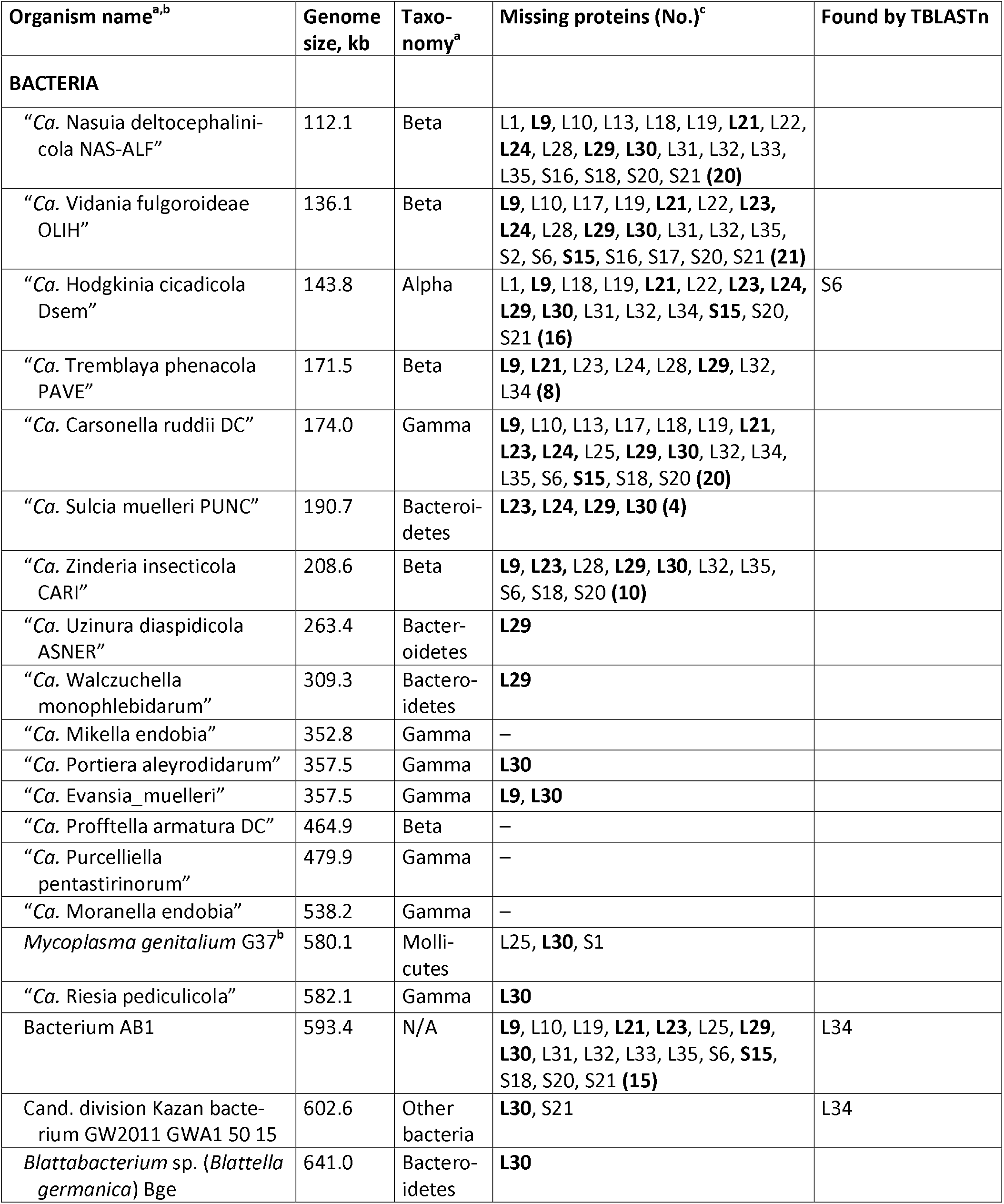
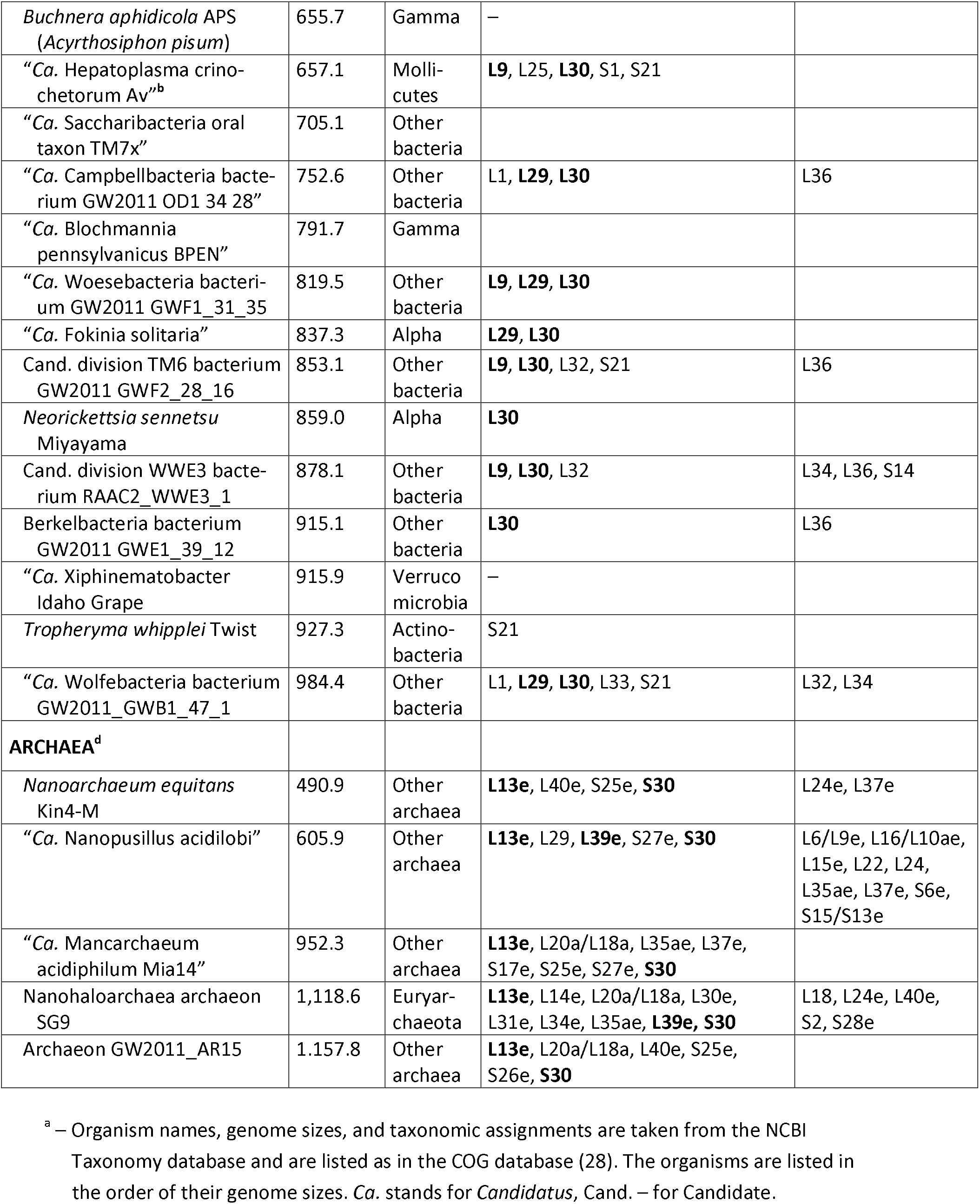
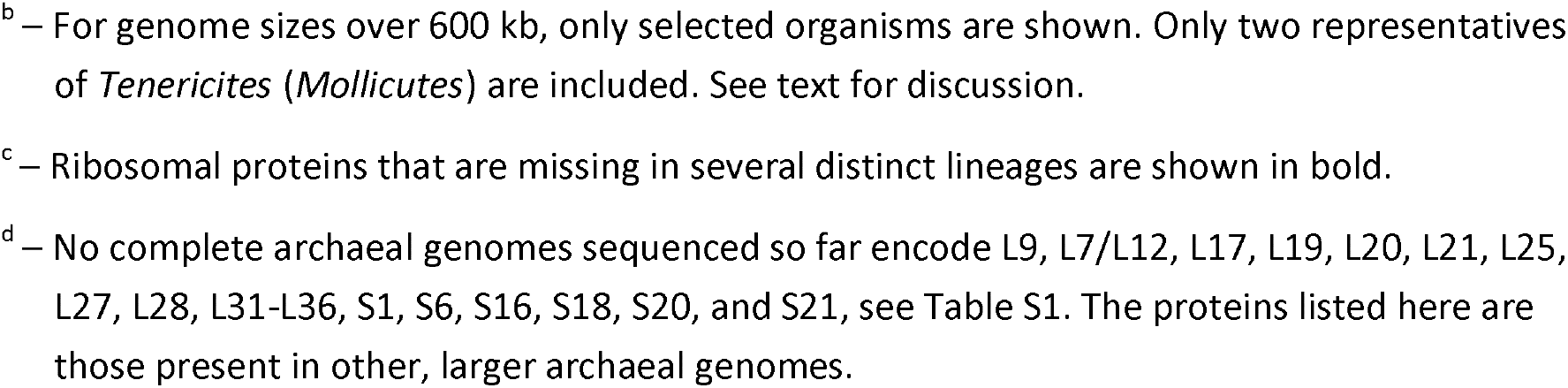
Ribosomal genes missing in organisms with tiny genomes.

### Loss of ribosomal protein genes in tiny genomes

Several previous studies investigated the gene contents in organisms with small genome sizes and reported a widespread absence of certain RP genes (2, 21, 25, 43). The most extensive loss of RP genes was observed in the tiny genomes of obligate insect symbionts that include members of *Alphaproteobacteria, Betaproteobacteria*, and *Bacteroidetes*. These genomes have undergone a dramatic compaction, resulting in genome sizes of less than 1.0 Mb and widespread loss of one or more RP genes (21, 25, 45). Indeed, in some of these tiny genomes, the loss of RP genes was extensive, such that up to 21 RP genes could be missing (Table 1). A massive loss of RP genes was also observed in the 593-kb genome of the bryozoan symbiont “bacterium AB1”, which is currently unclassified and apparently belongs to a novel major bacterial lineage (44).

However, not all bacteria with tiny genomes display massive loss of RP genes, and indeed, some of them retain nearly all RPs. The 263-kb genome of the flavobacterium “*Ca.* Uzinura diaspidicola”, an endosymbiont of armored scale insects, misses only a single RP gene, *rpmC*, that encodes L29 (Table 1). Similarly, the absence of *rpmC*, but no other RP gene, was observed in another flavobacterium, “*Ca.* Walczuchella monophlebidarum”, which has a slightly larger 309-kb genome. The 641-kb genome of yet another member of *Bacteroidetes, Blattabacterium* sp., also misses a single RP gene, in this case, the L30-encoding *rpmD*. The *rpmD* gene is also the only one missing in the genomes of the alphaproteobacterium *Neorickettsia sennetsu* (859 kb) and in some gammaproteobacteria, such as “*Ca.* Portiera aleyrodidarum” (357 kb), “*Ca.* Riesia pediculicola” (582 kb), and “*Ca.* Blochmannia pennsylvanicus” (792 kb). The 837-kb genome of “Ca. Fokinia solitaria”, an obligate intracellular endosymbiont of the ciliate *Paramecium* sp., lacks the genes for both L29 and L30 (Table 1).

Some tiny genomes actually encode the full set of core RPs (Figure 1A). In the investigated genome set, the smallest such genome (353 kb) was from the gammaproteobacterial symbiont of mealybugs “*Ca.* Mikella endobia”. This bacterium inhabits the cytoplasm of the betaproteobacterium “*Ca.* Tremblaya princeps”, which has an even smaller (171 kb) genome (45) and lacks the genes for eight RPs (Table 1). Other insect endosymbionts with tiny genomes that encode the full set of RPs include the alphaproteobacterial psyllid symbiont “*Ca.* Profftella armatura” (465 kb) and the gammaproteobacteria “*Ca.* Purcelliella pentastirinorum” and “*Ca.* Moranella endobia” (genome sizes, 480 kb and 539 kb, respectively, Table 2). The latter organism is also an intracellular symbiont of “*Ca.* Tremblaya princeps” (45).

**Table 2.** Lineage-specific loss of ribosomal proteins.

| Proteins (no.) | Lost in the following taxa (no. missing/no. organisms)<sup>a</sup> |
| --- | --- |
| **50S subunit** |  |
| L2-L6, L14- L16, L20 **(9)** | Always present |
| L7/L12, L10, L11, L27, L36 **(5)** | Missing only in poorly sequenced genomes |
| L9, L17- L19, L22, L23 **(6)** | Missing only in tiny genomes (Table 1) |
| L1, L13, L21, L24, L28 **(5)** | Missing in some tiny genomes (Table 1) and one additional (poorly sequenced?) genome |
| L25 | Missing in some tiny genomes (Table 1) and in *Coriobacteria* (11/11), *Bacillales* (6/50), *Lactobacillales* (14/23), *Mollicutes* (12/14), *Negativicutes* (10/10) |
| L29, L31, L32, L33 | Missing in tiny genomes (Table 1) and several other genome |
| L30 | Missing in some tiny genomes (Table 1), *Halanaerobiales* (6/6), *Pelagibacteriales* (2/2), *Rickettsiales* (5/9) |
| L34 | Missing in some tiny genomes (Table 1), *Halanaerobiales* (6/6), *Planctomycetes* (12/14) |
| L35 | Missing in some tiny genomes (Table 1), *Mollicutes* (2/14) |
| **30S subunit** |  |
| S1 | *Mollicutes* (11/14), *Dehalococcoidia* (2/2), *Erysipelothrix* (1/1) |
| S2 | Missing in two genomes: one tiny and one poorly sequenced |
| S3-S5, S8, S10, S11, S13, S14 **(8)** | Always present |
| S6, S15-S20 **(7)** | Missing only in tiny genomes, see Table 1 |
| S7, S9, S12 **(3)** | Missing in a single genome, possible sequencing error |
| S21 | *Actinobacteria* (155/155), *Deinococcus-Thermus* (6/6), *Fusobacteria* (6/6), *Halanaerobiales* (5/6), *Thermotogae* (9/9) |
| **Archaeal ribosomes** |  |
| L7ae, L12e, L15e, L18e, L19e, L24e, L32e, L37ae/L43a, L44e, S3ae, S4e, S6e, S8e, S19e, S24e, S28e **(16)** | Always present |
| L21e, L31e, L37e, S17e **(4)** | Missing in a single genome out of 122 |
| L40e, S27e | Missing in two genomes out of 122 |
| L13e | *Crenarchaeota* (12/25), *Euryarchaeota* (79/79), *Thaumarchaeota* (11/12) |
| L14e | *Archaeoglobi* (3/3), *Halobacteria* (31/31), *Methanomicrobia* (18/18), *Thermoplasmata* (10/10), *Thaumarchaeota* (11/12) |
| L20a/L18a | *Halobacteriales* (4/11), *Natrialbales* (5/11), *Thaumarchaeota* (12/12) |
| L30e | *Halobacteria* (31/31), *Thermoplasmata* (5/10) |
| L34e | *Archaeoglobi* (3/3), *Halobacteria* (31/31), *Methanomicrobia* (18/18), *Thermoplasmata* (10/10), *Thaumarchaeota* (12/12) |
| L35ae | *Archaeoglobi* (3/3), *Halobacteria* (31/31), *Methanomicrobia* (18/18), *Thermoplasmata* (10/10), *Thaumarchaeota* (12/12) |
| S25e, S26e, S30 | *Euryarchaeota* (79/79), tiny genomes |
| S27ae | *Haloferacales* (7/10) |

<sup>a</sup> – Data from the 1,309 genomes in the COG database (28); frameshifts and point mutations in RP genes (Table S3), as well as newly translated RPs (Table S4) are not counted as missing.

**Figure 1.**
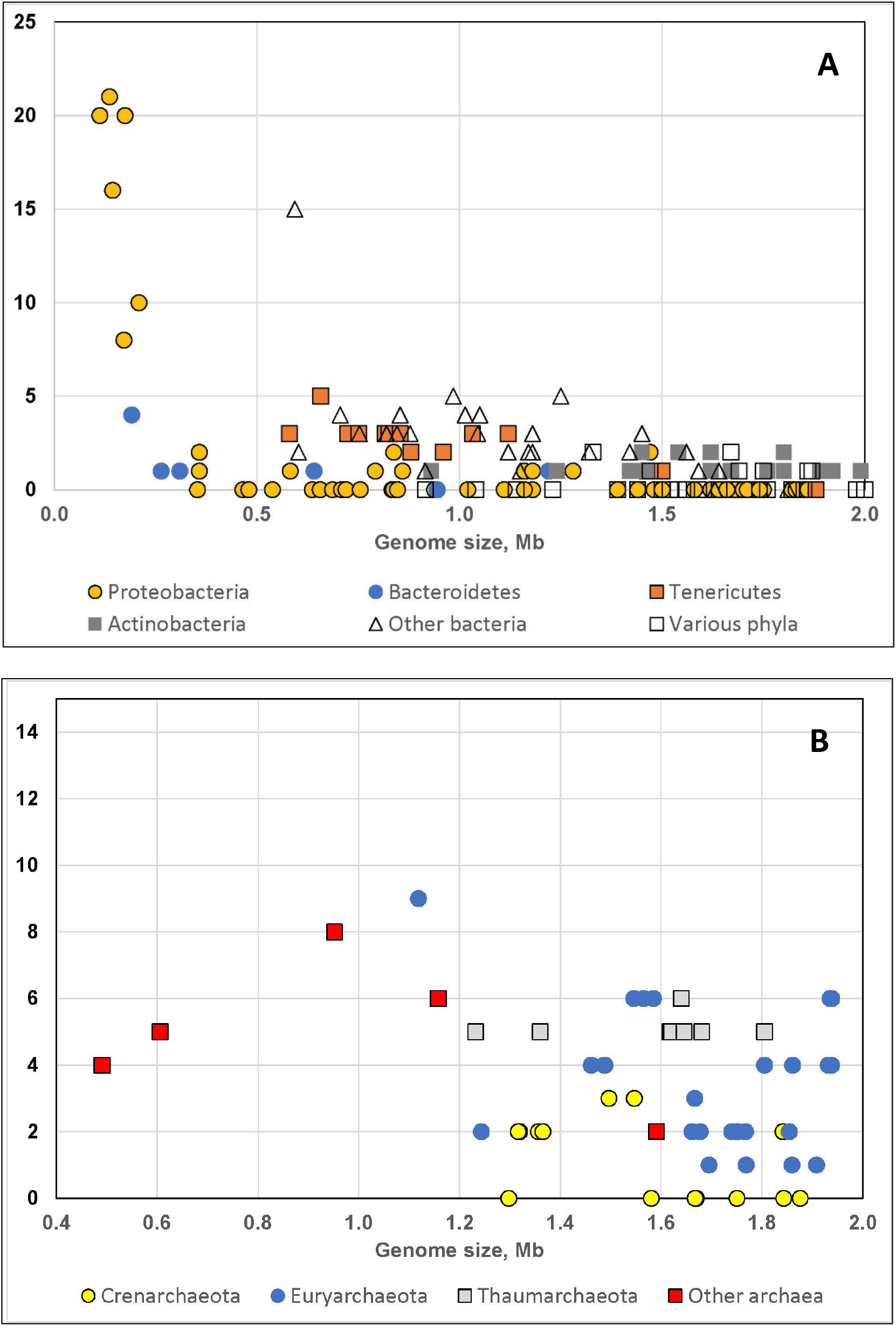
Loss of ribosomal genes in bacteria and archaea with small genome sizes. The number of ribosomal protein genes missing in various bacteria (A) and archaea (B) is shown as a function of the genome size. Each symbol indicates a representative organism from those included in the COG database (a single genome per genus). In A, yellow circles indicate the genomes of members of *Proteobacteria*, blue circles – *Bacteroidetes*, orange squares – *Tenericutes*, grey squares – *Actinobacteria*, empty squares - representatives of various phyla with 5-10 members in COGs (*Aquificae, Chlamydiae, Chloroflexi, Cyanobacteria, Fusobacteria, Spirochaetes, Synergistetes, Thermotogae*, and *Verrucomicrobia*); triangles indicate representatives of poorly sampled phyla (the “Other bacteria” group in COGs). In B, yellow circles indicate genomes of members of *Crenarchaeota*, blue circles – *Euryarchaeota*, grey squares – *Thaumarchaeota*, and red squares - representatives of poorly sampled phyla (the “Other archaea” group in COGs). See Table 1 for the names of representative organisms.

All 122 archaeal genomes included in the COG database lack 21 bacteria-specific RPs: L9, L7/L12, L17, L19-L21, L25, L27, L28, L31-L36, S1, S6, S16, S18, S20, and S21 (1, 4, 5), see Table S1. Only five of these 122 archaeal genomes are smaller than 1.2 Mb (Figure 1B); three of these small genomes come from the DPANN superphylum, one from Euryarchaeota, and one remains unclassified. These genomes show conservation of all the universal RPs and most archaea-specific RP genes. Each of these five genomes lacks the genes for L13e and S30, and in some of them, L20a/L18a and L39e genes are missing as well (Table 1). As mentioned above, a substantial number of RPs, nine in “*Ca.* Nanopusillus acidilobi” and five in “Nanohaloarchaea archaeon SG9”, are encoded in the respective genomes and are only missing in GenBank owing to the errors in genome submission (Tables 1 and S3).

### Lineage-specific loss of ribosomal protein genes

Figure 1A shows that, at genome sizes over 1.5 Mb, bacterial genomes rarely lack more than three RPs. At slightly larger genome sizes, most organisms contain the full RP sets. The exact position of the boundary between RP-missing and RP-complete protein sets varies between bacterial lineages but is typically around 2.0 Mb. The lowest such boundary, 0.8 Mb, was detected in *Gammaproteobacteria*: the only gammaproteobacterial genome in the COG that is larger than 0.8 Mb but is missing any RP genes is the above-mentioned genome of *Salinicola tamaricis*, where the absence of the *rpsB* gene is likely due to a sequencing error. In the analyzed genome set, the boundary for Betaproteobacteria and *Chloflexi* lies at 1.70 Mb, for *Bacteroidetes* at 1.88 Mb, for *Alphaproteobacteria* at 2.01 Mb, whereas for *Cyanobacteria* it is 3.34 Mb.

Irrespective of the genome size, no RP gene loss was observed in any representatives of the phyla *Aquificae* (9 genomes, 1.50 – 1.98 Mb), *Chlamydiae* (6 genomes, 1.04 – 3.07 Mb), *Chlorobi* (5 genomes, 2.15 – 3.29 Mb), *Spirochaetes* (11 genomes, 1.14 – 4.70 Mb), *Synergistetes* (5 genomes, 1.85 – 3.59 Mb), and the proteobacterial class *Epsilonproteobacteria* (12 genomes, 1.64 – 3.19 Mb), covered in the current version of COGs. Among poorly represented phyla (the “Other bacteria” group in COGs), the full set of RP genes was found in both members of *Armatimonadetes, Gemmatimonadetes*, and *Ignavibacteriae* and all three members of *Thermodesulfobacteria*. *Acidobacteria, Deltaproteobacteria*, and *Verrucomicrobia* had a single RP gene missing in a single organism, which could be due to the sequencing problems.

In certain lineages, however, loss of ribosomal genes was consistently detected in regular-sized genomes of free-living bacteria and archaea. As shown in Table 2, this type of RP gene loss is often lineage-specific. A striking example is the previously reported absence of the *rpsU* (S21) gene in every member of the phylum *Actinobacteria* (4). This trend still held true for the 155 actinobacterial genomes from 149 genera included in the current version of the COG database. An additional check in the NCBI protein database showed that S21 protein is not encoded in any genome from the phylum *Actinobacteria* sequenced to date. This protein is also missing in all representatives of the phyla *Deinococcus-Thermus, Fusobacteria*, and *Thermotogae* (Table 2, see also Table S3 in ref. (4)). All six representatives of the *phylum Fusobacteria* also lack the *rplY* (L25) gene, which is absent in certain lineages of *Actinobacteria, Firmicutes*, and *Tenericutes* as well (Table 2).

Similar lineage-specific patterns of gene loss were detected also in lower-level taxa. Thus, in the clostridial order *Halanaerobiales*, five of the six members: *Acetohalobium arabaticum, Halanaerobium hydrogeniformans, Halobacteroides halobius, Halocella* sp. SP3-1, and *Halothermothrix orenii*, with genomes in the 2.5 – 4.0 Mb range, lack *rpmD* (L30), *rpmH* (L34), and *rpsU* (S21) genes, whereas the remaining member, *Anoxybacter fermentans*, only lacks the first two. No other clostridial member in the COG system misses the *rpmH* or *rpsU* genes, pinpointing the loss of these genes to the base of the *Halanaerobiales* lineage. Likewise, in the order *Lactobacillales* (class *Bacilli*), the *rplY* (L25) gene is lost in members of three families, *Lactobacillaceae, Leuconostocaceae*, and *Streptococcaceae*, but present in the members of *Aerococcaceae, Carnobacteriaceae*, and *Enterococcaceae*.

Among the *Archaea*, the L30e protein is missing in all representatives of the euryarchaeal class *Halobacteria* and in all but one representative of the order *Thermoplasmatales* (Table 2).

### Widespread ribosomal protein gene loss in *Mollicutes*

The phylum *Tenericutes* presents a remarkable case of RP loss. Most early studies of gene essentiality focused on *Mycoplasma genitalium*, which has a 580-kb genome, the smallest among the known bacteria that are capable of axenic growth and can be obtained in pure culture (the recently sequenced genomes of several strains of *M. genitalium* are all at least 579.5 kb long) (18, 46). The genomes of *M. genitalium* and its close relative *Mycoplasma pneumoniae* were found to lack *rplY* (encoding L25 protein), *rpmD* (L30) and *rpsA* (S1) genes. Genes for all other core RPs were present and, with the possible exception of *rpmB* (L28), *rpsT* (S20) and *rpmGB* (encoding a paralog of L33), none could be disrupted by transposon mutagenesis (18, 22, 46).

Essentially the same pattern of the absence of the genes for L25, L30, and S1 has been detected in other mollicutes as well (21, 43). In the current version of the COGs, the coverage of this group was expanded to include representatives of 12 genera of *Mollicutes* and two recently sequenced unclassified members of the phylum *Tenericutes* (28). Among these 14 genomes, the genes for L25, L30, and S1 were missing, besides *Mycoplasma* spp., in representatives of four other genera: *Entomoplasma luminosum, Mesoplasma florum, Spiroplasma chrysopicola*, and *Ureaplasma parvum*. The genome of “*Ca.* Hepatoplasma crinochetorum”, in addition to these three genes, also lacked the genes for L9 and S21, whereas in four other mollicutes, L25 and S1 were missing; two of these genomes additionally lacked L35. Finally, the slightly larger (1.5-Mb) genome of *Acholeplasma laidlawii* only lacked L25, whereas the two unclassified members of the *Tenericutes*, “*Ca.* Izimaplasma str. HR1” and “Tenericutes bacterium MO-XQ” with their even larger genomes (1.88 and 2.16 Mb, respectively) were found to encode the full set of core RP genes.

### Experimentally identified non-essential ribosomal proteins

Over the past 15-20 years, numerous studies have been published aiming at the identification of the essential genes in a variety of bacteria (see Table S4). In the course of these projects, many genes, including certain RP-encoding genes, were identified as non-essential because their inactivation through transposon insertion or in-frame deletion proved to be non-lethal. We reviewed the relevant literature and compiled the lists of RP genes that have been successfully inactivated and therefore deemed non-essential (Table S4). Table S4 shows that the lists of non-essential RPs can vary dramatically between closely related organisms and, in some cases, even in experiments performed by different groups on the same bacterial strains. It should be noted that these lists include only the genes that have been explicitly reported to be disrupted. As an example, a detailed study of *Bacteroides thetaiotaomicron* strain VPI-5482 (47) identified only two dozen essential RP genes, suggesting that others could have been successfully inactivated. However, only mutants lacking L9 and L19 were used in subsequent experiments, positively marking these proteins as non-essential for *B. thetaiotaomicron*. Accordingly, numerous studies that centered on the essential genes and did not report the details of the disruption of non-essential genes [e.g. ref. (48) have been ignored. Nevertheless, a comparison of the data obtained on a variety of distinct organisms clearly shows that certain RPs are far more likely to be non-essential than the rest of the set. Furthermore, thorough analyses performed in *E. coli* and *B. subtilis* (6, 7, 49, 50) have resulted in closely similar lists of non-essential RPs (Tables S1 and S4).

### Loss propensity vs. non-essentiality of ribosomal proteins

Tables 1 and 2 show that certain RP genes are repeatedly identified as being prone to be lost in a variety of bacteria and archaea. Notably, some of the same genes could be successfully deleted in different organisms (Tables S1 and S4). Indeed, a comparison of the data in Tables 1, 2, and S4 reveals a consistent pattern: the genes that are often missing in tiny genomes are also non-essential in *E. coli* and/or *B. subtilis* (Table 3). Conversely, the genes that are always found even in tiny bacterial genomes could not be deleted from *E. coli* and, with the sole exception of L15, from B. subtilis (Table 3). Table S1 also shows that 12 genes that are dispensable in *E. coli* and/or *B. subtilis* are bacteria-specific, that is, missing in archaea and yeast. Indeed, the list of 26 dispensable RP genes (Table 3), includes 16 (of 21) bacteria-specific RPs and 10 (of 33) universal RPs. Thus, bacteria-specific RPs appear more likely to be non-essential than universal ones.

**Table 3.** Comparison of non-essentiality and gene loss of ribosomal proteins.

| Protein name<sup>a</sup> | Gene name | Deletion<sup>b</sup> |  | Loss in tiny genomes<sup>c</sup> |  |  |  |  |  |  |  | Lineage-specific gene loss<sup>d</sup> |
| --- | --- | --- | --- | --- | --- | --- | --- | --- | --- | --- | --- | --- |
|  |  | *E.coli* | *B.sub* | Nas | Vid | Hod | Tre | Car | Sul | Zin | AB1 |  |
| uL1 | RplA | Y | Y | Y | – | Y | – | – | – | – | – |  |
| bL9 | RplI | Y | Y | Y | Y | Y | Y | Y | – | Y | Y |  |
| uL10 | RplJ | – | – | Y | Y | – | – | Y | – | – | Y |  |
| uL11 | RplK | Y | Y | – | – | – | – | – | – | – | – |  |
| bL17 | RplQ | – | – | – | Y | – | – | Y | – | – | – |  |
| uL18 | RplR | – | – | Y | – | Y | – | Y | – | – | – |  |
| bL19 | RplS | – | – | Y | Y | Y | – | Y | – | – | Y |  |
| bL21 | RplU | – | – | Y | Y | Y | Y | Y | – | – | Y |  |
| uL22 | RplV | – | Y | Y | Y | Y | – | – | – | – | – |  |
| uL23 | RplW | – | Y | – | Y | Y | Y | Y | Y | Y | Y |  |
| uL24 | RplX | – | – | Y | Y | Y | Y | Y | Y | – | – |  |
| bL25 | RplY | Y | Y | – | – | – | – | Y | – | – | – | *Actinobacteria, Firmicutes* |
| bL28 | RpmB | – | Y | Y | Y | – | Y | – | – | Y | – |  |
| uL29 | RpmC | – | Y | Y | Y | Y | Y | Y | Y | Y | Y |  |
| uL30 | RpmD | – | – | Y | Y | Y | – | Y | Y | Y | Y | *Clostridia, Proteobacteria* |
| bL31 | RpmE | Y | Y | Y | Y | Y | – | – | – | – | Y | *Tenericutes* |
| bL32 | RpmF | Y | Y | Y | Y | Y | Y | Y | – | Y | Y |  |
| bL33 | RpmG | Y | Y | Y | – | – | – | – | – | – | Y |  |
| bL34 | RpmH | – | Y | – | – | Y | Y | Y | – | – | – | *Firmicutes, Planctomycetes* |
| bL35 | RpmI | Y | Y | Y | Y | – | – | Y | – | Y | Y | *Tenericutes* |
| bL36 | RpmJ | Y | Y | – | – | – | – | – | – | – | – |  |
| bS6 | RpsF | Y | Y | – | Y | – | – | Y | – | Y | Y |  |
| uS15 | RpsO | Y | – | – | Y | Y | – | Y | – | – | Y |  |
| bS18 | RpsR | – | – | Y | – | – | – | Y | – | Y | Y |  |
| bS20 | RpsT | Y | Y | Y | Y | Y | – | Y | – | Y | Y |  |
| bS21 | RpsU | Y | Y | Y | Y | Y | – | – | – | – | Y | *Actinobacteria, Deinococcus-Thermus, Fusobacteria, Thermotogae* |

<sup>a</sup> – Ribosomal protein names according to the universal nomenclature (5). Prefix ‘u’ indicates universal conservation of the protein, prefix ‘b’ indicates proteins that are specific for bacteria.
<sup>b</sup> – Successfully generated deletion mutants in *E. coli* (6, 53) and *B. subtilis* (7, 52) are indicated as Y, absence of such mutants is indicated by a dash. See Table S4 for details.
<sup>c</sup> – Organisms with tiny genomes are listed in the order of genome sizes (same as in Table 1): Nas, “*Ca. Nasuia* *deltoccephalinicola* str. NAS-ALF”; Vid, “*Ca. Vidania fulgoroideae* OLIH”; Hod, “*Ca. Hodgkinia cicadicola* Dsem”; Tre, “*Ca. Tremblaya phenacola* PAVE”; Car, “*Ca. Carsonella ruddii* DC”; Sul, “*Ca. Sulcia muelleri* PUNC”; Zin, “*Ca. Zinderia* *insecticola* CARI”; AB1, “bacterium AB1”. Y indicates the loss of the respective gene, presence of the gene is shown by a dash.
<sup>d</sup> – Bacterial phyla containing the lineages that exhibit loss of the respective genes (from Table 2).

We are unaware of systematic efforts on disruption or deletion of archaeal RPs. However, Table 2 shows that of the 33 archaea-specific RPs, 22 are conserved in nearly all analyzed genomes, whereas the rest exhibit lineage-specific gene losses.

### Loss of ribosomal protein genes in evolution vs ribosome structure and assembly

It is instructive to compare the pattern of RP loss during prokaryote evolution with the location of the respective RPs in the ribosome structure (51–56) and a related characteristic, the order of RP joining during the ribosome assembly (3, 57–61). Figure 2 and Table S5 show that neither of these features provides a clear-cut prediction of the RP loss propensity and/or (non)essentiality. Indeed, such frequently lost bacterial RPs as L9, L25, L29, L30, and S21 are located on the surface of the ribosome (Fig. 2A,B). However, L32, which is lost in many tiny genomes, has a significant buried area, whereas L34, which is lost in two bacterial lineages, is mostly buried in the ribosome structure. Conversely, several other surface RPs with relatively small buried areas (L16, L27, S16, S17, S18) are rarely lost and could not be deleted in either *E. coli* or *B. subtilis* (Tables 2 and 3). Likewise, most of the frequently lost RPs (L7/L12, L9, L25, L30, L32, S21, see Table S5) are incorporated into the ribosome at the late stages of its assembly (61). However, some of the RPs that join the ribosome early and interact with either 16S (S6, S20) or 23S (L21, L24, L29, L34) rRNA can also be lost or deleted (Table 3), whereas late-addition RPs L6, L16, L27, S2, S3, S10, S13, S14, and S19 are seldom lost (Table S5) and could not be deleted in *E. coli* or *B. subtilis*. Thus, there seems to be, at best, only a weak trend in the expected direction, namely, that RPs that are located on the surface of the ribosome and are attached late during the ribosome assembly are frequently lost in evolution and are often non-essential. A detailed accounting of specific protein-rRNA and protein-protein contacts (25) could eventually provide a better predictor of the RP loss propensity (non-essentiality).

**Figure 2.**
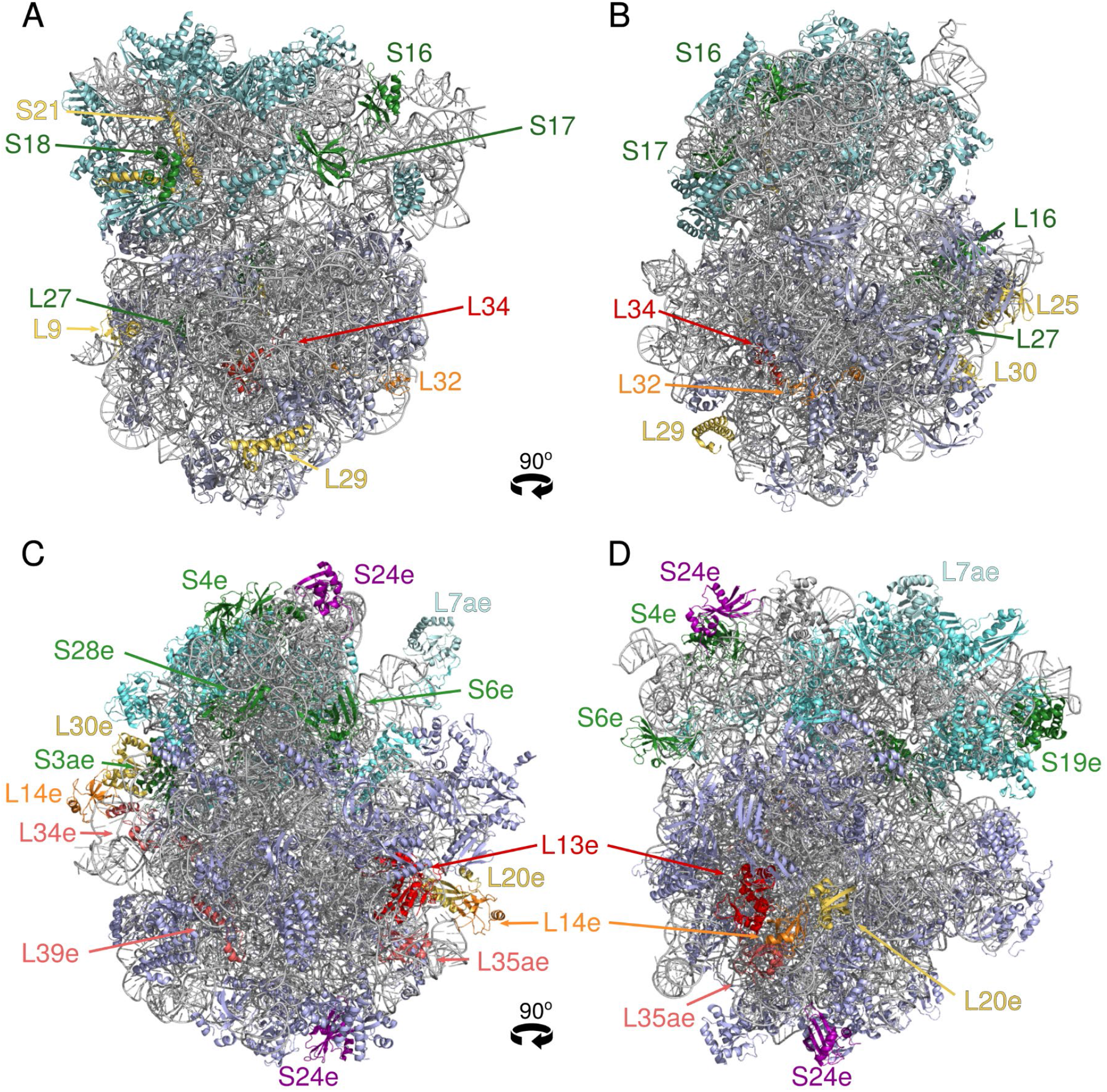
Localization of certain frequently and rarely lost surface proteins in the ribosomes of *Escherichia coli* and *Pyrococcus furiosus*. A, B. Crystal structure of the *E. coli* ribosome (PDB: 7K00), solved at 2 Å resolution by Watson *et al.*, 2020 (56). C, D. Cryo-EM structure of the *P. furiosus* ribosome (PDB: 4V6U), solved at 6.6 Å resolution by Armache *et al.*, 2012 (55). The ribosomal RNAs are shown in grey. Unless indicated otherwise, 50S subunit proteins are in wheat color and 30S subunit proteins are in cyan. Proteins mentioned in the text are indicated by bright colors: frequently lost surface proteins are in yellow, rarely lost ones are in green, other proteins described in the text are in red, orange and magenta. The structures were visualized and colored using PyMOL v. 1.0 (Schrödinger, LLC).

Similar trends are detectable among the archaea-specific RPs. The L13e protein, which is lost in all tiny archaeal genomes (Table 1) and is also missing in euryarchaea and nearly all thaumarchaea (Table 2), is only partially surface-exposed: its N-terminal loop and the first α-helix project deep into the core of the 50S subunit (Fig. 2C,D). Of the two ‘promiscuous’ surface proteins, L14e and S24e, that are present in two copies in the archaeal ribosome (55) (Fig. 2C,D), the former is often lost but the latter is never missing in archaea (Table 2). Among other frequently lost archaea-specific RPs (Table 2), L20a/L18a (aka LX) and L30e are surface proteins, but L34e and L35ae are mostly buried and L39e is only partially surface-exposed (54, 55). Conversely, surface-exposed proteins S3ae, S4e, S6e, S19e, and S28e (Fig. 2C,D) were never found to be missing in any of the analyzed archaeal genomes (Table 2).

## DISCUSSION

The overall conservation of the translation machinery among the bacteria, archaea, and eukaryotes (Table S1) is the strongest evidence of the common origin of all organisms, which allows including them in a single, universal Tree of Life (26, 27, 62). Indeed, 33 RPs are universal, that is, in all likelihood, have been conserved throughout the more than 3.5 billion years of the evolution of life (63, 64). Therefore, it is remarkable that genes for some of these universal RPs can be lost in many bacteria and archaea organisms with tiny genomes (Table 1) as well as in certain bacterial and archaeal lineages with larger genomes (Table 2), and can be deleted from the genomes of model bacteria without substantial loss of viability (Tables 3 and S4).

Here, we sought to trace the loss of RP genes in a relatively small and well-defined set of bacterial and archaeal genomes covered by the recent release of the COG database (28). This work was prompted by the observation that relatively few of the RP COGs had “perfect” phyletic patterns,that is, included representatives of all 1,309 organisms (or, in the case of domain-specific RPs, all representatives of either 1,187 bacteria or 122 archaea, respectively). Based on the previous studies (1–5, 21, 25, 43),, the missing RPs were expected to come primarily from the highly degraded genomes with an additional contribution of lineage-specific gene loss. These expectations proved to be largely correct, with organisms with tiny genomes (Table 1) and lineage-specific gene loss (Table 2) accounting for a large fraction of imperfect phyletic patterns among the RPs.

In addition, we identified multiple instances of frameshifted ORFs (Table S2) that were likely generated by sequencing errors. In certain cases, these frameshifts occurred in long stretches of identical nucleotides, which raises the possibility that some of them could represent authentic programmed frameshifts (40). This possibility, however, seems unlikely in cases where the genome of a closely related bacterium encodes an intact full-length ORF. Imperfect phyletic patterns can also be caused by problems in genome annotation whereby certain ORFs, particularly short ones, are overlooked by the annotation software (Table S3). Given the widespread loss of RP genes (Tables 1 and 2), it would not be realistic to require every newly sequenced genome to contain the full set of RP genes. Nevertheless, when a bacterial or archaeal genome of more than 1 Mb in size lacks any of the 43 widely conserved RPs (Table 2), it should raise a red flag. Further, the example of “*Ca.* Nanopusillus acidilobi” (Table 1) shows that short and/or divergent RPs from poorly studied bacteria and archaea should not be deemed pseudogenes without clear evidence that this is indeed the case. In particular, as shown in Table 2, almost all archaeal genomes encode the 33 universal and 20 archaea-specific RPs, so that the absence of any of these genes in an archaeal genome is highly unlikely.

So, what conclusions can be drawn from the patterns of RP loss – and conservation – patterns shown in Tables 1, 2, and 3? First, these observations validate the previously noted trend of independent loss of orthologous RP genes in several phylogenetically distant lineages (4). Examples include the loss of L25 in certain members of *Actinobacteria, Firmicutes*, and *Mollicutes*, loss of L30 in some members of *Clostridia* and *Alphaproteobacteria*, loss of L34 in certain *Clostridia* and *Planctomycetes*, and the loss of S21 in several distinct bacterial phyla (Table 2). Among archaea-specific RPs, it is worth noting the simultaneous absence of L13e, L14e, L20a, L34e, and L35ae proteins in many members of *Euryarchaeota* and *Thaumarchaeota* (Table 2).

The second prominent trend is the gradual loss of RPs within a single lineage. Thus, previous analyses of the mollicute genomes reported the absence of the genes for L25, L30, and S1 (21, 43). This pattern was confirmed here for several mollicute genomes, albeit not for the two recently sequenced unclassified members of *Tenericutes*. These comparisons allowed us to reconstruct a possible scenario of RP gene loss in the phylum *Tenericutes* (Figure 3). It appears that progressive genome reduction during the evolution of the *Mollicutes* first led to the loss of L25, then S1, followed by either L30 or L35, and culminated in the loss of two more genes in “*Ca.* Hepatoplasma crinochetorum” (Fig. 3). Remarkably, the same set of genes coding for L25, L30, and S1 that is missing in Mycoplasma spp. is also missing in the genome of *Erysipelothrix* rhusiopathiae, a member of the Firmicutes branch that is closest to the *Mollicutes* (65).

**Figure 3.**
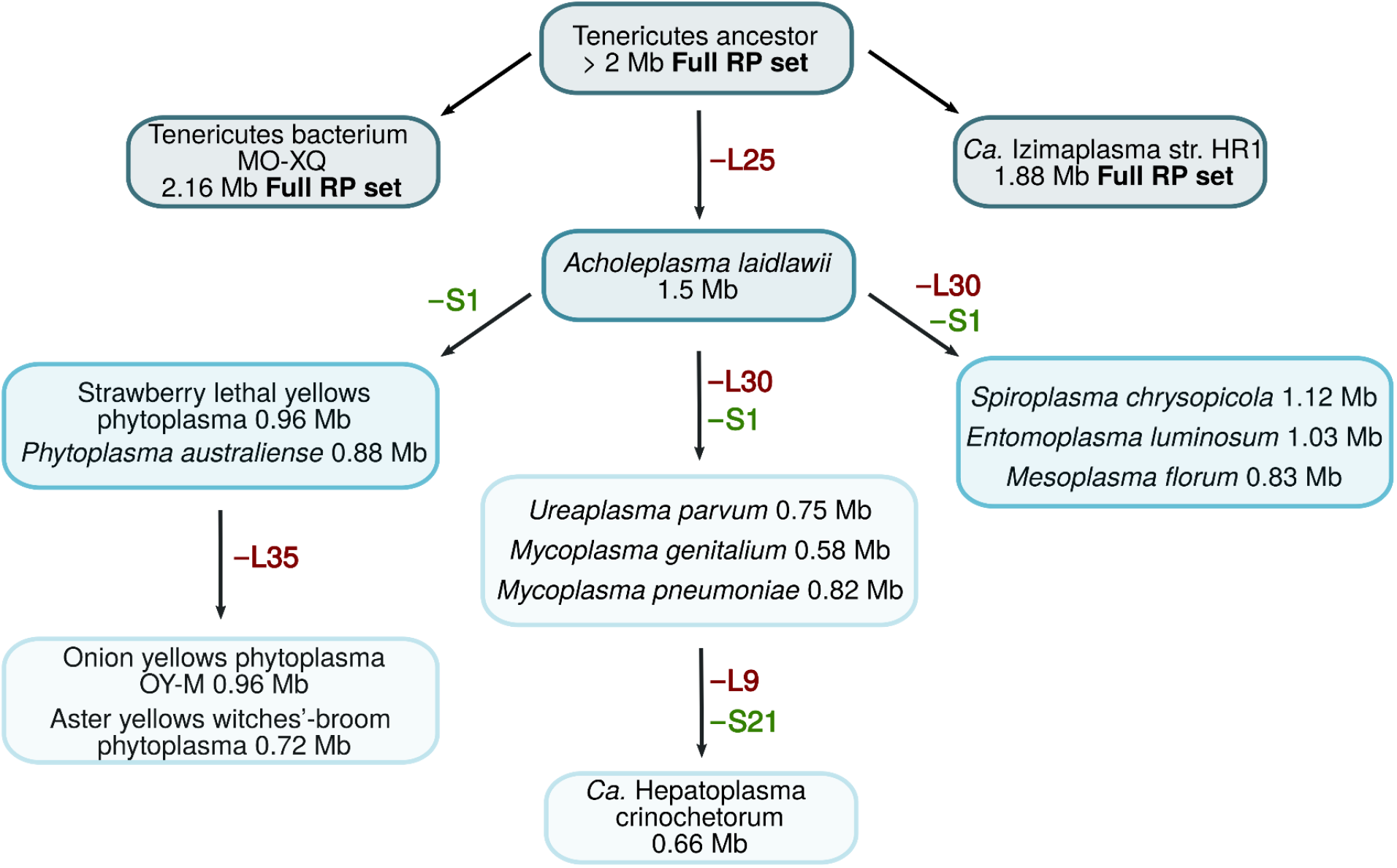
A possible scenario of ribosomal protein loss in the *Mollicutes* (*Tenericutes*). The organisms are those listed in the COG database. Based on the phylogeny of this group, which divides *Mollicutes* into at least four distinct lineages, *Acholeplasma/Phytoplasma*, *Spiroplasma/ Mesoplasma, Mycoplasma pneumoniae/Ureaplasma*, and *Mycoplasma hominis* (76), the loss of L30 and S1 may have occurred independently on two or more occasions. The loss of of L25 may have occurred at an early stage in the evolution of this group or has occurred several times.

The loss of RPs in phylum- or class-level lineages generally correlates with two other hallmarks of non-essentiality, the availability of deletion mutants in model organisms and frequency of loss in tiny genomes (Table 3). The genes that encode apparently non-essential RPs, but for which few or no losses were observed in the genomes included in the COGs, are likely to be lost in other bacterial or archaeal genomes, especially, small ones. For example, the loss of L1 and/or L9, which was detected in several tiny genomes from “Other bacteria” (Table 1), has been reported to be widespread among the Candidate Phyla Radiation (Patescibacteria), a vast and diversified group of poorly characterized bacteria that are thought to be symbionts or parasites of other bacteria (66).

An interesting aspect of the non-essential RP gene set is its potential use in synthetic biology. In previous attempts to construct a “minimal” bacterial cell, either fully synthetic (23, 67–69) or highly streamlined (70), the researchers aimed at obtaining rapidly growing microorganisms and chose not to tweak their RP gene content. Accordingly, the synthetic *Mycoplasma genitalium* JCVI-1.0 (GenBank accession number CP000925) and both synthetic versions of *Mycoplasma mycoides*, JCVI-syn1.0 (GenBank: CP002027) and JCVI-syn3.0 (GenBank: CP016816), included all 50 RP genes that are normally found in these organisms (which do not encode L25, L30, and S1). The synthetic genome of *Caulobacter ethensis*-2.0 included all core RPs of *Caulobacter crescentus* except for L1 and L6 (69). The MiniBacillus project ended up including all 54 core RP genes (Table S1), as well as YlxQ (L7ae) and paralogs of L6, L33, and S14 (70, 71). Future attempts at constructing streamlined bacterial genomes might involve attempts to substantially reduce the sets of RP genes. In contrast, in archaea, the overall conservation of the RPs leaves few choices for such gene deletion.

It should be noted that the absence of RP genes was discussed here – and elsewhere – in terms of genome compaction and lineage-specific gene loss, based on the presence of the respective RP genes in the genomes of closely related organisms. However, a recent study (72) has shown that certain RPs are encoded by phages, which indicated the distinct possibility of the acquisition of the RP genes through lateral transfer. The L7/L12 and S21 genes were most widespread in phages, and some phages also encoded L9 and S30. Further, analysis of viral metagenomes revealed the occasional presence of genes for L11, L19, L31, L33, S6, S9, S15, and S20 and, in some instances, for L2 and L10 (72). The presence of such genes in phage genomes could be explained by the pressure on the phage to provide the cell with its own RPs to accelerate translation, particularly when the respective genes are missing in the host genome. Indeed, the RPs listed above are often lost, both in organisms with tiny genomes and in specific bacterial lineages (Tables 1 and 2).

Overall, the observations presented here show that the evolution of RPs is more malleable and dynamic than previously thought. It remains to be seen whether additional massive sequencing of diverse bacterial and archaeal genomes leads to further erosion of the set of universal RPs and/or of those that are conserved within the archeal or bacterial domains of life.

## MATERIALS AND METHODS

### Genome coverage and protein selection

The list of bacterial and archaeal genomes used in this work was taken from the recent release of the COG database (28). This set includes 1,309 complete genomes of 1,187 bacteria and 122 archaea, most of them with a single representative of the respective genus (1,234 named genera, see https://ftp.ncbi.nih.gov/pub/COG/COG2020/data/cog-20.org.csv for the full list).

The list of RPs analyzed in this work was also taken from the COG database (the ‘Ribosome 30S subunit’, ‘Ribosome 50S subunit’, and ‘Archaeal ribosomal proteins’ groups in the ‘COG Pathways’ list, https://www.ncbi.nlm.nih.gov/research/cog/pathways). This list included 54 bacterial (or universal) proteins and 33 archaea-specific proteins, as listed in the Supplementary Table S1. Two auxiliary RPs, S22 (RpsV, Sra) and S31e (Thx), were not included in this survey because there were no respective COGs in the database. S22 protein is mostly expressed during the stationary phase and appears to be non-essential for the viability of *E. coli* (38). S31e (Thx) is part of the 30S subunit in *Thermus thermophilus* (53) and is mostly found in Bacteroidetes, Proteobacteria, and several other phyla. The RNA-binding protein L7ae (YlxQ) is associated with archaeal ribosomes but apparently not with bacterial ribosomes (73). The S1 protein was deemed present when the respective ORF included three or more S1-like domains. The archaeal protein set did not include L38e and L41e proteins (arCOG04057 and arCOG06624 in (74), respectively), which are not represented in the current set of COGs.

### Identification of missing ribosomal genes

The list of RPs missing from each genome was taken from the phyletic profiles of the respective COGs. The nucleotide sequences of the respective genomes were searched with representative RP sequences (taken either from Table S1 or from closely related taxa) using the recent version of the TBLASTn program (41) that allows the selection of specific organisms based on the NCBI Taxonomy assignments. The resulting BLAST hits (cut-off E-value, 0.1) were verified using the CDD-search (75) and compared against the protein sets in GenBank and RefSeq databases. The confirmed RP ORFs that were missing in GenBank were classified as either frameshifted (or, in several cases, those missing a recognizable start codon) or overlooked; for the latter ones, the full-size ORFs were translated from the genomic sequences using ORFfinder (https://www.ncbi.nlm.nih.gov/orffinder/). Representative frameshifted and overlooked ORFs are listed, respectively, in Table S2 and Table S3.

The RPs that produced no statistically significant hits in TBLASTn searches were classified as missing in the respective genomes. These genomes were classified into tiny (less than 1 Mb in size for bacteria or 1.2 Mb for archaea) and regular-sized; they were further sorted by phyla according to their COG assignments (which rely on the NCBI Taxonomy database).

### Identification of non-essential ribosomal genes

The lists of non-essential ribosomal genes (Table S4) were collected from the literature and two online databases, Database of Essential Genes [DEG, (13)] and the Online Gene Essentiality database [OGEE, (14)]. Since these databases, and most of the original literature, focused on essential genes, the supplementary material files for each paper were individually checked to select those genes that had been positively identified as non-essential and ignore those genes that were not listed as essential but whose status had not been specified.

To assess the ribosomal localization of selected RPs, the structures of ribosomes of *E. coli* (PDB: 7K00) (56) and *Pyrococcus furiosus* (PDB: 4V6U) (55) were downloaded from the Protein Data Bank and visualized using PyMOL v. 1.0 (Schrödinger, LLC). Individual surface proteins were colored based on their loss propensity (Table 2).

## ACKNOWLEDGEMENTS

This study was supported by the Intramural Research Program of the U.S. National Library of Medicine at the National Institutes of Health.

## Supplementary Tables

**Table S1. Core bacterial and archaeal proteins, their unified nomenclature of Ban *et al.* 2014, Pfam domains, and the UniProt entries in *Escherichia coli, Bacillus subtilis, Mycoplasma pneumoniae, Aeropyrum pernix, Haloarcula marismortui* and *Saccharomyces cerevisiae*.**

**Table S2. Frameshifted ribosomal proteins in the genomes covered by the COG database**

**Table S3. Unannotated ORFs coding for ribosomal proteins in the genomes covered by the COG database**

**Table S4. Experimental data on large-scale inactivation of ribosomal proteins**

**Table S5. Loss of ribosomal proteins that differ in their rRNA interactions and the order of assembly, as described by Chen and Williamson, 2013.**

