## Supplemental Tables S1 to S5 for "Non-essential ribosomal proteins in bacteria and archaea identified using COGs"

### Supplementary Tables

**Table S1. Core bacterial and archaeal proteins, their unified nomenclature from Ban *et al*., 2014 (1), Pfam (2)entries, and the UniProt (3) accession numbers in *Escherichia coli*, *Bacillus subtilis*, *Mycoplasma pneumoniae*, *Aeropyrum pernix*, *Haloarcula marismortui* and *Saccharomyces cerevisiae*.**

**Table S2. Frameshifted ribosomal proteins in the genomes covered by the COG database**

**Table S3. Unannotated ORFs coding for ribosomal proteins in the genomes covered by the COG database**

**Table S4. Experimental data on large-scale inactivation of ribosomal proteins**

**Table S5. Loss of ribosomal proteins that differ in their rRNA interactions and the order of assembly**

**Table S1. Core bacterial and archaeal proteins, their unified nomenclature, and the UniProt and Pfam entries in *Escherichia coli*, *Bacillus subtilis*, *Mycoplasma pneumoniae*, *Aeropyrum pernix*, *Haloarcula marismortui* and *Saccharomyces cerevisiae*[a].**

| Protein name | Gene name | COG number[b] | Pfam entry[b] | Ban, 2014[c] name | Distribution[d] | | UniProt accession number[e] | | | | |
|---|---|---|---|---|---|---|---|---|---|---|---|
| | | | | | Ban, 2014[d] | COG | *E. coli* | *B.subtilis* | *Halo-arcula* | *Aero-pyrum* | Yeast |
| **50S subunit** | | | | | | | | | | | |
| L1 | RplA | COG0081 | PF00687 | uL1 | BAE | A+B | P0A7L0 | Q06797 | P12738 | Q9Y9W6 | P0CX44 |
| L2 | RplB | COG0090 | PF00181+ PF03947 | uL2 | BAE | A+B | P60422 | P42919 | P20276 | Q9YFN1 | P0CX45 |
| L3 | RplC | COG0087 | PF00297 | uL3 | BAE | A+B | P60438 | P42920 | P20279 | Q9YFM2 | P14126 |
| L4 | RplD | COG0088 | PF00573 | uL4 | BAE | A+B | P60723 | P42921 | P12735 | Q9YFM1 | P10664 |
| L5 | RplE | COG0094 | PF00281 | uL5 | BAE | A+B | P62399 | P12877 | P14124 | Q9YF87 | P0C0W9 |
| L6 | RplF | COG0097 | PF00347 | uL6 | BAE | A+B | P0AG55 | P46898 | P14135 | Q9YF91 | P05738 |
| L9 | RplI | COG0359 | PF03948 | bL9 | B | B | P0A7R1 | P37437 | – | – | – |
| L10 | RplJ | COG0244 | PF00466 | uL10 | BAE | A+B | P0A7J3 | P42923 | P15825 | Q9Y9W8 | P05317 |
| L11 | RplK | COG0080 | PF03946+ PF00298 | uL11 | BAE | A+B | P0A7J7 | Q06796 | P14122 | Q9Y9W5 | P0CX53 |
| L7/L12 | RplL | COG0222 | PF16320+ PF00542 | bL12 | B | B | P0A7K2 | P02394 | – | – | – |
| L13 | RplM | COG0102 | PF00572 | uL13 | BAE | A+B | P0AA10 | P70974 | P29198 | Q9YB50 | P26784 |
| L14 | RplN | COG0093 | PF00238 | uL14 | BAE | A+B | P0ADY3 | P12875 | P22450 | Q9YF82 | P0CX41 |
| L15 | RplO | COG0200 | PF00828 | uL15 | BAE | A+B | P02413 | P19946 | P12737 | Q9YF98 | P02406 |
| L16/ L10AE | RplP | COG0197 | PF00252 | uL16 | BAE | A+B | P0ADY7 | P14577 | P60617 | Q9YEY5 | P41805 |
| L17 | RplQ | COG0203 | PF01196 | bL17 | B | B | P0AG44 | P20277 | – | – | – |
| L18 | RplR | COG0256 | PF00861 | uL18 | BAE | A+B | P0C018 | P46899 | P14123 | Q9YF94 | P26321 |
| L19 | RplS | COG0335 | PF01245 | bL19 | B | B | P0A7K6 | O31742 | – | – | – |
| L20 | RplT | COG0292 | PF00453 | bL20 | B | B | P0A7L3 | P55873 | – | – | – |
| L21 | RplU | COG0261 | PF00829 | bL21 | B | B | P0AG48 | P26908 | – | – | – |
| L22 | RplV | COG0091 | PF00237 | uL22 | BAE | A+B | P61175 | P42060 | P10970 | Q9YF76 | P05740 |
| L23 | RplW | COG0089 | PF00276 | uL23 | BAE | A+B | P0ADZ0 | P42924 | P12732 | Q9YFM0 | P04456 |
| L24 | RplX | COG0198 | PF00467 | uL24 | BAE | A+B | P60624 | P0CI78 | P10972 | Q9YF83 | P05743 |

| | | | | | | | | | | |
|---|---|---|---|---|---|---|---|---|---|---|
| L25 | RplY | COG1825 | PF01386 | bL25 | B | B | P68919 | A0A6M3Z BP3 | – | – | – |
| L27 | RpmA | COG0211 | PF01016 | bL27 | B | B | P0A7L8 | P05657 | – | – | – |
| L28 | RpmB | COG0227 | PF00830 | bL28 | B | B | P0A7M2 | P37807 | – | – | – |
| L29 | RpmC | COG0255 | PF00831 | uL29 | BAE | A+B | P0A7M6 | P12873 | P10971 | P58085 | P0CX84 |
| L30/L7E | RpmD | COG1841 | PF00327 | uL30 | BAE | A+B | P0AG51 | P19947 | P14121 | Q9YF96 | P05737 |
| L31 | RpmE | COG0254 | PF01197 | bL31 | B | B | P0A7M9 | Q03223 | – | – | – |
| L32 | RpmF | COG0333 | PF01783 | bL32 | B | B | P0A7N4 | O34687 | – | – | – |
| L33 | RpmG | COG0267 | PF00471 | bL33 | B | B | P0A7N9 | P56849 | – | – | – |
| L34 | RpmH | COG0230 | PF00468 | bL34 | B | B | P0A7P5 | P05647 | – | – | – |
| L35 | RpmI | COG0291 | PF01632 | bL35 | B | B | P0A7Q1 | P55874 | – | – | – |
| L36 | RpmJ | COG0257 | PF00444 | bL36 | B | B | P0A7Q6 | P20278 | – | – | – |
| **30S subunit** | | | | | | | | | | |
| S1 | RpsA | COG0539 | PF00575 | bS1 | B | B | P0AG67 | P38494 | – | – | – |
| S2 | RpsB | COG0052 | PF00318 | uS2 | BAE | A+B | P0A7V0 | P21464 | P29202 | Q9YB45 | P32905 |
| S3 | RpsC | COG0092 | PF07650+ PF00189 | uS3 | BAE | A+B | P0A7V3 | P21465 | P20281 | Q9YF78 | P05750 |
| S4 | RpsD | COG0522 | PF00163+ PF01479 | uS4 | BAE | A+B | P0A7V8 | P21466 | Q00862 | Q9YB58 | O13516 |
| S5 | RpsE | COG0098 | PF00333+ PF03719 | uS5 | BAE | A+B | P0A7W1 | P21467 | P26815 | Q9YF95 | P25443 |
| S6 | RpsF | COG0360 | PF01250 | bS6 | B | B | P02358 | P21468 | – | – | – |
| S7 | RpsG | COG0049 | PF00177 | uS7 | BAE | A+B | P02359 | P21469 | P32552 | Q9YAU8 | P26783 |
| S8 | RpsH | COG0096 | PF00410 | uS8 | BAE | A+B | P0A7W7 | P12879 | P12742 | Q9YF89 | P0C0W1 |
| S9 | RpsI | COG0103 | PF00380 | uS9 | BAE | A+B | P0A7X3 | P21470 | P05763 | Q9YB48 | P0CX51 |
| S10 | RpsJ | COG0051 | PF00338 | uS10 | BAE | A+B | P0A7R5 | P21471 | P23357 | Q9YAV2 | P38701 |
| S11 | RpsK | COG0100 | PF00411 | uS11 | BAE | A+B | P0A7R9 | P04969 | P10788 | Q9YB55 | P06367 |
| S12 | RpsL | COG0048 | PF00164 | uS12 | BAE | A+B | P0A7S3 | P21472 | Q5UZR8 | Q9YAU5 | P0CX29 |
| S13 | RpsM | COG0099 | PF00416 | uS13 | BAE | A+B | P0A7S9 | P20282 | Q00861 | Q9YB60 | P0CX55 |
| S14 | RpsN | COG0199 | PF00253 | uS14 | BAE | A+B | P0AG59 | P12878 | P26816 | P58731 | P41057 |
| S15 | RpsO | COG0184 | PF00312 | uS15 | BAE | A+B | P0ADZ4 | P21473 | P05762 | Q9YCX3 | P05756 |
| S16 | RpsP | COG0228 | PF00886 | bS16 | B | B | P0A7T3 | P21474 | – | – | – |
| S17 | RpsQ | COG0186 | PF00366 | uS17 | BAE | A+B | P0AG63 | P12874 | P12741 | Q9YF81 | P0CX47 |
| S18 | RpsR | COG0238 | PF01084 | bS18 | B | B | P0A7T7 | P21475 | – | – | – |

| | | | | | | | | | | | |
|---|---|---|---|---|---|---|---|---|---|---|---|
| S19 | RpsS | COG0185 | PF00203 | uS19 | BAE | A+B | P0A7U3 | P21476 | P20284 | Q9YF74 | Q01855 |
| S20 | RpsT | COG0268 | PF01649 | bS20 | B | B | P0A7U7 | P21477 | – | – | – |
| S21 | RpsU | COG0828 | PF01165 | bS21 | B | B | P68681 | P21478 | – | – | – |
| **Archaeal ribosomal proteins** | | | | | | | | | | | |
| L7Ae | Rpl7Ae | COG1358 | PF01248 | eL8 | AE | A+B | – | P46350 | P12743 | Q9YAX7 | P48589 |
| L12E/L44/L45/RPP1/RPP2 | RPP1A | COG2058 | PF00428 | P1/P2 | AE | A | – | – | P15772 | Q9Y9W9 | P05318 |
| L13E | RPL13 | COG4352 | PF01294 | eL13 | AE | A | – | – | P29198 | Q9YEN9 | P40212 |
| L14E/L6E/L27E | RPL14A | COG2163 | PF01777 | eL14 | AE | A | – | – | – | Q9YDD7 | P0C2H6 |
| L15E | RPL15A | COG1632 | PF00827 | eL15 | AE | A | – | – | P60618 | Q9YBZ8 | P05748 |
| L18E | RPL18A | COG1727 | PF17135 | eL18 | AE | A | – | – | P12733 | Q9YB51 | P0CX49 |
| L19E | RPL19A | COG2147 | PF01280 | eL19 | AE | A | – | – | P14119 | Q9YF93 | P0CX82 |
| L20A (L18A, LX) | RPL20A RPL18ae RplX | COG2157 | PF01775 | eL20 | E | A | – | – | P14125 | P58289 | P0CX23 |
| L21E | RPL21A | COG2139 | PF01157 | eL21 | AE | A | – | – | P12734 | P58077 | Q02753 |
| L24E | RPL24A | COG2075 | PF01246 | eL24 | AE | A | – | – | P14116 | Q9Y9A7 | P04449 |
| L30E | RPL30E | COG1911 | PF01248 | eL30 | AE | A | – | – | – | Q9YAX7 | P14120 |
| L31E | RPL31A | COG2097 | PF01198 | eL31 | AE | A | – | – | P18138 | Q9YD25 | P0C2H8 |
| L32E | Rpl32e | COG1717 | PF01655 | eL32 | AE | A | – | – | P12736 | Q9YF92 | P38061 |
| L34E | RPL34A | COG2174 | PF01199 | eL34 | AE | A | – | – | – | P58026 | P87262 |
| L35AE/L33A | Rpl35A | COG2451 | PF01247 | eL33 | AE | A | – | – | – | Q9Y9G4 | P05744 |
| L37E | RPL37A | COG2126 | PF01907 | eL37 | AE | A | – | – | P32410 | Q9YEQ4 | P49166 |
| L39E | RPL39 | COG2167 | PF00832 | eL39 | AE | A | – | – | P22452 | P59472 | P04650 |
| L40E | RPL40A | COG1552 | PF01020 | eL40 | AE | A | – | – | Q5UYU5 | Q9YFY7 | P0CH08 |
| L44E | RPL42A | COG1631 | PF00935 | eL42 | AE | A | – | – | P32411 | Q9YF00 | P0CX27 |
| L37AE/L43A | RPL43A | COG1997 | PF01780 | eL43 | AE | A | – | – | P60619 | Q9YC06 | P0CX25 |
| S3AE | RPS3A | COG1890 | PF01015 | eS1 | AE | A | – | – | Q5V296 | Q9YCV8 | P33442 |

| | | | | | | | | | | | |
|---|---|---|---|---|---|---|---|---|---|---|---|
| S4E | RPS4A | COG1471 | PF00900+ PF16121 | eS4 | AE | A | – | – | P22510 | Q9YF85 | P0CX35 |
| S6E (S10) | RPS6A | COG2125 | PF01092 | eS6 | AE | A | – | – | P21509 | Q9Y9B6 | P0CX37 |
| S8E | RPS8A | COG2007 | PF01201 | eS8 | AE | A | – | – | P49402 | Q9YDY0 | P0CX39 |
| S17E | RPS17A | COG1383 | PF00833 | eS17 | AE | A | – | – | Q5V5R5 | Q9YA67 | P02407 |
| S19E (S16A) | RPS19A | COG2238 | PF01090 | eS19 | AE | A | – | – | P19952 | Q9YD22 | P07280 |
| S24E | RPS24A | COG2004 | PF01282 | eS24 | AE | A | – | – | P19953 | Q9YCY0 | P0CX31 |
| S25e | RPS25 | COG4901 | PF03297 | eS25 | AE | A | – | – | – | Q9Y914 | Q3E792 |
| S26e | RPS26B | COG4830 | PF01283 | eS26 | E | A | – | – | – | Q05DX2 | P39938 |
| S27AE | RPS27ae | COG1998 | PF01599 | eS31 | AE | A | – | – | Q06125 | P61297 | P05759 |
| S27E | RPS27A | COG2051 | PF01667 | eS27 | AE | A | – | – | Q5UX21 | Q9YF01 | P35997 |
| S28E/ S33 | RPS28A | COG2053 | PF01200 | eS28 | AE | A | – | – | A0A4P8 K2X9 | Q9Y9A6 | Q3E7X9 |
| S30 | RPS30 | COG4919 | PF04758 | eS30 | AE | A | – | – | – | Q9Y9T9 | P0CX33 |

[a] – A more detailed version of this table is available in Excel format as Supplementary Table S1A.

[b] – These COG and Pfam numbers can be used to view the respective database entries, e.g., for L1 protein, as https://www.ncbi.nlm.nih.gov/research/cog/cog/COG0081/ and http://pfam.xfam.org/family/PF00687.

[c] – Ribosomal protein names in the universal nomenclature of Ban *et al*., 2014 (1).

[d] – Presence of the respective proteins in bacteria (B), archaea (A) and eukaryotes (E) according to Ban *et al*. (2014) and in COGs. Discrepancies between the two are highlighted yellow.

[e] – These accession numbers can be used to view the respective protein entries in the NCBI protein database and in UniProt, e.g. for L1 protein, as https://www.ncbi.nlm.nih.gov/protein/P0A7L0 and https://www.uniprot.org/uniprot/P0A7L0 (*E. coli*), https://www.ncbi.nlm.nih.gov/protein/Q06797 and https://www.uniprot.org/uniprot/Q06797 (*B. subtilis*), and so on. Green shading indicates *E.coli* proteins with viable full-length deletion mutants in the Keio collection (4) and those *B. subtilis* proteins that are marked non-essential in SubtiWiki (5).

**Table S2. Frameshifts and point mutations in ribosomal protein genes in the genomes covered by the COG database**

| Organism name, genome GenBank accession no. | Genome size, Mb | Taxonomy in COGs | Pro-tein | COG | Frame 1 boundaries, protein ID (if available) | Frame 2 boundaries, protein ID (if available) |
|---|---|---|---|---|---|---|
| *Actinobaculum* sp. 313, CP029033.1 | 2.77 | Actino-bacteria | L2 | COG0090 | Frame +1: 857650..858100 | Frame +3: 858099..858464 |
| | | | L14 | COG0093 | Frame +2: 861053..861328 | Frame +1: 861334..861417 |
| *Advenella kashmirensis* WT001, AFK64574.1 | 4.42 | Beta | L1 | COG0081 | Frame -2: 4318292..4318771 | Frame -1: 4318095..4318293 |
| | | | L6 | COG0097 | Frame -2: 4158914..4159123 | Frame -1: 4158609..4158920 |
| | | | L21 | COG0261 | Frame -2: 1061612..1061875 | Frame -1: 1061574..1061615 |
| | | | S6 | COG0360 | Frame -1: 2704266..2704538 | Frame -3: 2704189..2704242 |
| *Anaerotignum propionicum* DSM 1682, CP014223.1 | 3.12 | Clostridia | S4 | COG0522 | Frame +2: 585926..586072 | Frame +1: 586075..586461 |
| *Atlantibacter hermannii* NCTC12129, LR134136.1 | 4.55 | Gamma | L23 | COG0089 | Frame +1: 428506..428781; VDZ71398.1 | Frame +3: 428748..428789 |
| | | | S9 | COG0103 | Frame +1: 504925..504996 | Frame +2: 504992..505315 |
| *Azorhizobium caulinodans* ORS 571, AP009384.1 | 5.37 | Alpha | L9 | COG0359 | Frame -3: 4895567..4895716; BAF90303.1 | Frame -2: 4895154..4895636; BAF90302.1 |
| *Breoghania* sp. L-A4, CP031841.1 | 5.03 | Alpha | S5 | COG0098 | Frame +1: 3228187..3228483 | Frame -3: 3227942..3228184 |
| *Brucella melitensis* str. 16M AE008917.1, AE008917.1 | 3.29 | Alpha | L27 | COG0211 | Frame -2: 209218..209487, wrong translation start in AAL51384.1 | |
| "*Ca.* Atelocyanobacterium thalassa" isolate ALOHA, CP001842.1 | 1.44 | Cyano-bacteria | S17 | COG0186 | Frame +3: 1181154..1181309 | Frame +1: 1181314..1181391 |
| "Ca. Methanomethylophilus alvus" Mx-05, CP017686.1 | 1.67 | Eury-archaeota | S14 | COG0199 | Frame -3: 1401381 to 1401527, no N-terminal fragment | |
| "*Ca.* Nanopusillus acidilobi", CP010514.1 | 0.61 | Other archaea | S14 | COG0199 | Frame -3: 604224..604298 | Frame -1: 604160..604207 |
| "*Ca.* Tenderia electrophaga" CP013099 | 3.76 | Gamma | S19 | COG0185 | Frame +3: 877839..879123, 94 aa, no start codon | |
| "*Ca.* Uzinura diaspidicola" ASNER, CP003263.1 | 0.26 | Bactero-idetes | L21 | COG0261 | Frame +1: 100210..100434 | Frame +3: 100422..100508 |

| | | | | | | |
|---|---|---|---|---|---|---|
| *Desulfotalea psychrophila* LSv54, CR522870.1 | 3.66 | Delta | L6 | COG0097 | Frame +2; 1279586..1279756, CAG35868.1 | Frame +2: 1279755..1280131, CAG35869.1 |
| | | | S8 | COG0096 | Frame +3: 1279164..1279361 | Frame +2: 1279352..1279561 |
| *Enterococcus faecalis* V583, AE016830.1 | 3.36 | Bacilli | L13 | COG0102 | Frame -3: 3100862..3100966 | Frame -1: 3100543..3100860 |
| *Faecalibacterium prausnitzii* A2165, CP022479.1 | 3.11 | Clostridia | L22 | COG0091 | Frame +1: 2112610..2112750 | Frame +3: 2112726..2112941 |
| *Frondihabitans* sp. PAMC 28766, CP014513.1 | 4.77 | Actino-bacteria | L7/L12 | COG0222 | Frame -3: 3920472..3920576 | Frame -2: 3920254..3920415 |
| | | | S8 | COG0096 | Frame +3: 1279164..1279361 | Frame +2: 1279352..1279561 |
| *Gottschalkia acidurici* 9a, CP003326.1 | 3.11 | Tissierellia | L9 | COG0359 | Frame -2: 3004298..3004453, | Frame -1: 3004155..3004271 |
| *Ketogulonicigenium vulgare* Y25, CP002224.1 | 3.29 | Alpha | S18 | COG0238 | Frame -3: 1093683..1093817 | Frame -1: 1093583..1093654 |
| *Lactobacillus acidophilus* NCFM, CP000033.3 | 1.99 | Bacilli | S19 | COG0185 | Frame +3: 293403..293528 | Frame +1: 293482..293685 |
| *Laribacter hongkongensis* HLHK9, CP001154.1 | 3.17 | Beta | S9 | COG0103 | Frame -2: 2660379..2660627 | Frame -3: 2660237..2660416 |
| *Massilia putida* 6NM-7T, CP019038.1 | 7.52 | Beta | L11 | COG0080 | Frame +3: 4335990..4336055 | Frame +2: 4336052..4336417 |
| *Melioribacter roseus* P3M-2, CP003557.1 | 3.3 | Other bacteria | S4 | COG0522 | Frame -1: 232066..232254 | Frame -3: 231632..232075 |
| *Methylomusa anaerophila* MMFC1, AP018449.1 | 4.78 | Negati-vicutes | L35 | COG0291 | Frame +1: 3868123..3868230, BBB92833.1 | Frame +2: 386226.. 3868318 |
| Onion yellows phytoplasma OY-M, AP006628.2 | 0.85 | Mollicutes | L3 | COG0087 | Frame +1: 243267..243902, BAD04285.1 Lost start codon results in 65 N-terminal aa missing | |
| | | | L24 | COG0198 | Frame +3: 248499..248693 | Frame +1: 248680..248838 |
| | | | S4 | COG0522 | Frame -3: 658421..658540; BAD04671.1 | Frame +2: 657948..658430 |
| *Paenalcaligenes hominis* 15S00501, CP019697.1 | 2.69 | Beta | L2 | COG0090 | Frame -1: 1598924..1599670 | Frame -3: 1598847..1598924 |
| | | | L3 | COG0087 | Frame -3: 1601258..1600791 | Frame -2: 1600585.. 1600788 |
| *Parabacteroides distasonis* ATCC 8503, CP000140.1 | 4.81 | Bactero-idetes | S4 | COG0522 | Frame -3: 2777237..2777734 | Frame -2: 2777133..2777264 |
| *Parachlamydia acanthamo-ebae* UV-7, FR872580.1 | 3.07 | Chlamy-diae | S15 | COG0184 | Frame -1: 1022709..1022873 | Frame -2: 1022606..1022704 |

| | | | | | | |
|---|---|---|---|---|---|---|
| *Paraglaciecola psychrophila* 170 , CP003837.1 | 5.41 | Gamma | L2 | COG0090 | Frame -3: 4969102..4969206 | Frame -2: 4968386..4969063, AGH47270.1 |
| | | | L31 | COG0254 | Frame +2: 218429..218527 | Frame +1: 218527..218619, AGH42341.1 |
| *Plesiomonas shigelloides* MS-17-188, CP027852.1 | 3.97 | Gamma | S8 | COG0096 | Frame -2: 2081499..208159 | Frame -1: 2081212..2081493 |
| *Providencia stuartii* MRSN 2154 | 4.4 | Gamma | L7/L12 | COG0222 | Frame -2: 1846220.. 1846558, AFH93584.1 | Frame -3: 1846193.. 1846227 |
| *Rodentibacter pneumotropi-cus* NCTC8284, LR134405.1 | 2.44 | Gamma | L7/L12 | COG0222 | Frame +3: 225381..225467 | Frame +2: 225530..225745 |
| | | | S16 | COG0228 | Frame -3: 2228925..2228972 | Frame -2: 2228728..2228937 |
| | | | S20 | COG0268 | Frame +3: 2158104..2158226 | Frame +1: 2158279..2158371 |
| | | | L27 | COG0211 | Frame +1: 575761..576006, no start codon, point mutation | |
| *Salinicola tamaricis* F01, CP023559.1 | 4.28 | Gamma | L6P | COG0097 | Frame -2: 1172703..1172332 | Frame -1: 1172314..1172180 |
| | | | L7/L12 | COG0222 | Frame -3: 1193430..1193516 | Frame -2: 1193182..1193373<br>Frame -1: 1193153..1193194 |
| | | | L9 | COG0359 | Frame -1: 742979..743326 | Frame -3: 742917..742979 |
| | | | L25 | COG1825 | Frame -1: 3431492..3431671 | Frame -3: 3431097..3431501 |
| | | | S4 | COG0522 | Frame -3: 1167843..1168277 | Frame -2: 1167665..1167730 |
| *Simkania negevensis* Z, FR872582.1 | 2.63 | Chlamy-diae | L31 | COG0254 | Frame +1: 787441..787494 | Frame +2: 787496..787678, CCB88630.1 |
| *Sulfodiicoccus acidiphilus* HS-1, AP018553.1 | 2.35 | Cren-archaeota | L2 | COG0090 | Frame +1: 267295..267639, BBD71897.1 | Frame: +3: 267636..268019, BBD71898.1 |
| *Sulfuricella denitrificans* skB26, AP013066.1 | 3.22 | Beta | L9 | COG0359 | Frame -2: 1812117..1812246, BAN35601.1 | Frame -1: 1811804..1812166, BAN35600.1 |
| *Syntrophus aciditrophicus* SB CP000252.1, | 3.18 | Delta | S12 | COG0048 | Frame +2: 311162..311347 | Frame +3: 311373..311525 |
| *Thioalkalivibrio nitratiredu-cens* DSM 14787, CP003989.2 | 4.00 | Gamma | L36 | COG0257 | Frame -2: 2648149..2648093 | Frame -3: 2648085..2648038 |
| *Thioclava nitratireducens* 25B10_4, CP019437.1 | 4.24 | Alpha | S16 | COG0228 | Frame -3: 611211..611582, 123 aa, no start codon | |
| *Xanthobacter autotrophicus* Py2, CP000781.1 | 5.63 | Alpha | L14 | COG0093 | Frame +1: 1902232..1902666, ABS66934.1 | Frame +2: 5308729..5308830 (not adjacent) |

**Table S3. Examples of unannotated ORFs coding for ribosomal proteins in the genomes covered by the COG database**

| Organism name, GenBank accession no. | Genome size, Mb | Taxonomy in COGs | Ribosomal protein | COG no. | Newly translated ORF; locus tag, if available |
|---|---|---|---|---|---|
| **BACTERIA** | | | | | |
| *Sulfobacillus acidophilus* TPY, CP002901.1 | 3.55 | Clostridia | L27 | COG0211 | ORF: 607200..607481 Frame: +3; 93 aa |
| | | | L28 | COG0227 | ORF: 1069175..1069366, Frame: -1; 63 aa |
| | | | L32 | COG0333 | ORF: 1133208..1133387, Frame: +3; 58 aa |
| | | | L33 | COG0267 | ORF: 306096..306302, Frame: +3; 68 aa |
| | | | L36 | COG0257 | ORF: 343563..343673, Frame: = +3; 37 aa |
| | | | S14 | COG0199 | ORF: 337812..337997, Frame: +3 ; 61 aa |
| *Pelotomaculum thermopropionicum* SI, AP009389.1 | 3.03 | Clostridia | L28 | COG0227 | ORF: 1841595..1841786, Frame: +3; 63 aa |
| | | | L32 | COG0333 | ORF: 1830440..1830619, Frame: -1; 59 aa |
| | | | L34 | COG0230 | ORF: 3024909..3025082, Frame: -3; 57 aa |
| | | | S14 | COG0199 | ORF: 335426..335614, Frame: +2; 62 aa |
| | | | S21 | COG0828 | ORF: 901026..901331, Frame: +3; 101 aa |
| *Herpetosiphon aurantiacus* DSM 785, CP000875.1 | 6.79 | Chloroflexi | L28 | COG0227 | ORF: 438793..438975, Frame: +1; 60 aa |
| | | | L29 | COG0255 | ORF: 6258620.. 6258832, Frame: +2; 70 aa |
| | | | L35 | COG0291 | ORF: 5927538..5927750, Frame: -2; 70 aa |
| | | | L36 | COG0257 | ORF: 6266198..6266311, Frame: +2; 38 aa |
| *"Ca.* Methylomirabilis oxyfera", FP565575.1 | 2.75 | Other bacteria | L28 | COG0227 | ORF: 1181939..1182139, Frame: +2; 67 aa |
| | | | L33 | COG0267 | ORF:459805..459951, Frame: +1; 49 aa |
| | | | L34 | COG0230 | ORF: 2546125..2546259, Frame: +1; 44 aa |
| | | | L36 | COG0257 | ORF: 488708..488818, Frame: +2; 37 aa |
| | | | S13 | COG0099 | ORF: 488935..489300, Frame: +1, 124 aa |
| "Candidate division WWE3 bacterium RAAC2 WWE3 1", CP006914.1 | 0.88 | Other bacteria | L34 | COG0230 | ORF: 795925..796083, Frame: -1; 52 aa |
| | | | L36 | COG0257 | ORF: 890953..890843, Frame: -2; 38 aa |
| | | | S14 | COG0199 | ORF: 73005..73229, Frame: +3; 74 aa |
| *Planktomarina temperata* RCA23, CP003984.1 | 3.29 | Alphaproteo bacteria | L34 | COG0230 | ORF: 2522654..2522788, Frame: +2; 44 aa |
| | | | L36 | COG0257 | ORF: 3024835..3024960, Frame: = +1; 41 aa |
| *"Ca.* Paracaedimonas acanthamoebae", CP008936.1 | 2.18 | Alphaproteo bacteria | L9 | COG0359 | ORF: 611520 to 611951, Frame: +3; 144 aa |
| | | | L32 | COG0333 | ORF: 164521..164676, Frame: -3; 59 aa |
| | | | L36 | COG0257 | ORF: 1218708..1218833, Frame: +3; 41 aa |

| | | | | | |
|---|---|---|---|---|---|
| *Rugosibacter aromatici-vorans* Ca6, CP010554.1 | 2.93 | Betaproteo-bacteria | L36 | COG0257 | ORF: 2557159..2557269, Frame: -3; 37 aa |
| | | | S14 | COG0199 | ORF: 2561406..2561747, Frame: -1; 113 aa |
| "Candidatus Wolfebacteria bacterium GW2011 GWB1 47 1", CP011209.1 | 0.98 | Other bacteria | L32 | COG0333 | ORF: 333695..333901, Frame: -3; 68 aa |
| | | | L34 | COG0230 | ORF: 178951..179097, Frame: -1; 48 aa |
| *"Ca.* Tenderia electrophaga" CP013099.1 | 3.76 | Gammapro-teobacteria | L36 | COG0257 | ORF: 885686..885796, Frame: +2; 37 aa |
| | | | L30/L7E | COG1841 | ORF: 883593..883778, Frame: +3; 61 aa |
| *Hydrogenimonas* sp. MAG, AP019005.1 | 2.19 | Epsilonpro-teobacteria | L36 | COG0257 | ORF: 2048969..2048859, Frame: -2; 37 aa |
| | | | S14 | COG0199 | ORF: 2053957..2054142, Frame: -1; 61 aa |
| *Paucimonas lemoignei* NCTC10937, LS483371.1 | 5.92 | Betaproteo-bacteria | L36 | COG0257 | ORF: 5452130..5452246, Frame: -3; 38 aa |
| | | | S13 | COG0099 | ORF: 5451643..5451999, Frame: -1; 118 aa |
| | | | S12 | COG0048 | ORF: 5466930..5467298, Frame: -2; 123 aa |
| *Salinicola tamaricis* F01, CP023559.1 | 4.28 | Gammapro-teobacteria | L4 | COG0088 | ORF 1178149..1178727, Frame: -3; 192 aa |
| | | | S18 | COG0238 | ORF: 744224..744472, Frame: -1; 82 aa |
| **ARCHAEA** | | | | | |
| *"Ca.* Nanopusillus acidilobi", CP010514 | 0.61 | Other archaea | L6P/L9E | COG0097 | ORF: 501349..501882, Frame: -2; 177 aa; locus tag: Nps_02895 |
| | | | L15e | COG1632 | ORF: 250345..250866, Frame: +1; 173 aa; locus tag: Nps_01385 |
| | | | L16/L10AE | COG0197 | ORF 579086..579643, Frame: -1; 185 aa; locus tag: Nps_03305 |
| | | | L22 | COG0091 | ORF: 586054..586629,( Frame: +1; 191 aa; locus_tag: Nps_03365 |
| | | | L24 | COG0198 | ORF: 503210..50374, Frame: -1, 174 aa; locus_tag: Nps_02910 |
| | | | S6e | COG2125 | ORF: 329344..329733; Frame -2; 129 aa; locus tag: Nps_01880 |
| | | | S15P/S13E | COG0184 | ORF: 269039..269431, Frame: -1; 130 aa; locus tag: Nps_01520 |
| | | | L35ae | COG2451 | ORF: 558084..558428, Frame -3; 114 aa; locus tag: Nps_03205 |
| | | | L37e | COG2126 | ORF: 580869..581003, Frame -3; 54 aa |
| *"Ca.* Nitrosocosmicus oleophilus", CP012850.1 | 3.43 | Thaum-archaeota | L24e | COG2075 | ORF: 865559..865771, Frame +2, 70 aa |

| | | | | | |
|---|---|---|---|---|---|
| *Nanoarchaeum equitans* Kin4-M, AE017199.1 | 0.49 | Other archaea | L24e | COG2075 | ORF: 297703..297867, Frame +1; 54 aa |
| | | | L37e | COG2126 | ORF: 5537..5707, Frame: -1; 56 aa |
| “Nanohaloarchaea archaeon SG9”, CP012986.1 | 1.12 | Eury-archaeota | L18 | COG0256 | ORF: 47533..48027, Frame: -1; 169 aa; locus tag: AQV86_00305 |
| | | | S2 | COG0052 | ORF: 413675..414298, Frame: -3; 207 aa; locus tag: AQV86_02305 |
| | | | S28e | COG2053 | ORF: 411826..411990, Frame: -1; 58 aa |
| | | | L24e | COG2075 | ORF: 411666..411818, Frame +2, 50 aa |
| | | | L40e | COG1552 | ORF: 708355..708516, Frame +1, 53 aa |
| *Thermofilum adornatus*, CP006646.1 | 1.75 | Cren-archaeota | L34e | COG2174 | ORF: 785414..785677, Frame -2, 87 aa |

**Table S4. Experimental data on large-scale inactivation of ribosomal protein-coding genes**

| **Organism** | **Inactivated or deleted genes**[a] | **Reference** |
|---|---|---|
| *Acinetobacter baumannii* | **L9**, **L19**, L22, L27, **L31**, **L32**, **L33**, **S1**, **S20** | (6) |
| *Acinetobacter baylyi* | **L9**, L27, **L31**, **L33**, L36 | (7) |
| *Agrobacterium fabrum* | **L9**, L11, L15, **L31**, **L32**, **L33**, S7, S12, S17, S19, S21 | (8) |
| *Bacillus subtilis* | L11, **L25**, **S1** | (9) |
| | **L1**, **L9**, L11, **L15**, **L22**, L23, **L25**, **L28**, **L29,** **L31**, **L32**, **L33**, L34, **L35**, **L36**, **S6**, **S20**, **S21** | (10, 11) |
| | **L1**, L23, **L34**, **L36**, **S6** | (12) |
| *Bacteroides fragilis* | **L9**, L17, **L19**, **L32**, L34 | (13) |
| *Bacteroides thetaiotaomicron* | **L9**, **L19** | (14) |
| *Brevundimonas subvibrioides* | **L9**, L30, **L31**, **L35**, S15, S16 | (8) |
| *Burkholderia cenocepacia* | **L9**, *L18*, **L19**, L21, **L25**, *L30*, *L36*, *S2*, *S13*, S15, S17, *S18*, ***S21*** | (15, 16) |
| *Burkholderia thailandensis* | **L1, L9**, L21, **L31**, S2, S3, S15 | (17) |
| *Campylobacter jejuni* | **L9**, L20, **L25**, **L33**, **L35**, S14 | (18) |
| | L3, **L9**, L21, **L33**, **S1**, S2, S4, S5, S7, S17 | (19) |
| | ***L9***, **L19**, L20, *L27*, ***L31***, **S1**, *S4*, *S8*, S15, S18 | (20) |
| *Caulobacter crescentus* | **L1,** L10, **L28**, **L29,** L30, **L31**, **L33**, **L35** | (21) |
| *Escherichia coli* | L5, **L9**, L16, **L22**, **S1**, S3, S7 | (22) |
| | **L1**, **L9**, L11, **L25**, **L31**, **L32**, **L33**, **L35**, **L36**, **S6**, S15, **S20**, **S21** | (4) |
| | **S6**, S9, S13, S15, S17, **S20** | (23) |
| | **L15**, L21, L24, L27, **L29,** L30, **L34**, S9, S17 | (24) |
| | **L1, L9**, **L31**, **L32**, **L33**, **S6**, S9, S15, S18, **S20** (*L13, S1, S2, S4, S7, S12, S17, S18*) | (25) |
| | **L31**, L36, **S1**, S15, S18 | (26) |
| | **L1, L9**, **L11**, **L25**, **L31**, **L32**, **L33**, **L35**, **L36**, **S6**, S15, **S20**, **S21** | (27) |
| *Francisella novicida* | **L9**, **L19**, **L33**, **S1**, **S21** | (28) |
| *Francisella tularensis* | **L1, S1** | (29) |
| *Haemophilus influenzae* | L7, **L9** | (30) |
| *Helicobacter pylori* | L10, L18, **L19**, **L33**, S11, S13, S16 | (31) |
| *Mycobacterium avium* | L7/L12, **L9**, L10, **L19**, **L25**, **L28**, **L31**, **L32**, **L33**, **S6**, S11, S14, S15, S17, S18 | (32). |
| *Mycobacterium tuberculosis* | **L1,** L15, **L28**, **L33**, L36, S14, S16, S18 | (33) |
| | **L9**, L15, **L25**, **L28**, L30, **L33**, S14, S16, S18 | (34) |
| | **L1,** L5, **L9**, L22, **L25**, **S1**, S2 | (35) |
| *Mycoplasma bovis* | L34 | (36) |
| *Mycoplasma pulmonis* | **L28**, **L33**, **S1**8 | (37, 38) |

| | | |
|---|---|---|
| *Neisseria gonorrhoeae* | L24, L27, L30, **L31**, **L32**, **L33**, L36, S10, **S21** | (39) |
| *Porphyromonas gingivalis* | **L9**, L17, **L19**, **L31**, **L32**, L34, **S1**, S16, S18, **S21** | (40) |
| *Providencia stuartii* | **L19**, **L31**, **S1** | (41) |
| *Pseudomonas aeruginosa* | L3, L4, **L9**, **L19**, **L25**, **L32**, S3, S4, S5,S12 | (42) as cited in (7) |
| | L2, L3, L4, **L9**, **L19**, L21, L23, **L25**, S4, S5, **S6**, S8, S9, S12, **S21** | (43). |
| | **L9**, L13, L17, L21, **L25**, L27, **L28**, **L31**, **L33**, L34, **S1**, **S6**, S9, S10, S15, S18, **S20** | (44) |
| | **L9**, **L25**, **L31**, **L32**, **L33**, S15 | (45) |
| | **L9**, **L25**, **L31**, **L32**, **L33**, S2, **S21** | (46) |
| *Pseudomonas protegens* | **L9**, **L25**, **L31**, **L33**, L36 | (47) |
| *Rhodopseudomonas palustris* | **L9**, L21, **L31**, **L33**, **S1**, S2, S16 | (48) |
| *Rubrivivax gelatinosus* | **L9**, **L25**, **L33** | (49) |
| *Salmonella enterica* | L16 | (50) |
| | **L1, L9**, **L31**, **L32**, **L33**, L36, **S20** | (51) |
| | **L1, L9**, L10, **L25**, **L31**, **L32**, **L33**, L36, **S1**, S9, S15, **S20**, **S21** | (52) |
| *Shewanella oneidensis* MR-1 | **L19** | (53) |
| *Staphylococcus aureus* | L6, S17 | (54) |
| | **L19**, **L25**, L27, **L28**, **L33**, **S1**, S14 | (55) |
| | **L9**, **L33**, **S20** | (56) |
| | **L1, L9**, L11, L21, L23, **L25**, **L28**, **L32**, **L33**, L34, **S1**, **S20** | (57) |
| *Streptococcus agalactiae* | **L9**, **L28**, S2, **S6**, S14, **S20** | (58) |
| *Streptococcus mutans* | **L9**, **L19**, **L28**, **L32**, **L33**, **L35**, **S1**, **S20**, **S21** | (59) |
| *Streptococcus pyogenes* | L7/L12, **L9**, L10, L15, L23, **L28**, **L29, L32**, S14, S16, **S20** | (60) |
| *Streptococcus sanguinis* | **L9**, L15, L21, L24, **L28**, **L29,** L30, **L31**, **L32**, **L33**, **L35**, L36, S9, S13, S15, **S20**, **S21** | (61) |
| *Streptococcus suis* | **L9**, L10, **L28**, **L32**, **L33**, **L35**, **S1**, S2, S12, **S20** | (62) |
| *Synechococcus elongatus* | **L9**, L15, **L28** | (63) |
| *Vibrio cholerae* | **L25**, **L31**, **L33** | (64) |
| | **L9**, **L31**, **L32** | (65) |
| Candidate phyla radiation | **L1, L9**, L30 | (66) |

[a] – The proteins whose genes that could be inactivated in several distinct lineages are shown in bold. Those proteins for which the data varied between strains, are shown in italics.

**Table S5. Loss of ribosomal proteins that differ in their rRNA interactions and the order of assembly**

| 50S subunit protein | Assembly order[a] | Missing in genomes[b] | | 30S subunit protein | Assembly order[a] | Missing in genomes[b] |
|---|---|---|---|---|---|---|
| L20 | I | 0 | | S4 | I | 0 |
| L21 | I | 7 | | S6 | I | 4 |
| L22 | I | 3 | | S8 | I | 0 |
| L24 | I | 7 | | S15 | I | 3 |
| L1 | II | 5 | | S16 | I | 2 |
| L3 | II | 0 | | S17 | I | 1 |
| L4 | II | 0 | | S18 | I | 3 |
| L13 | II | 3 | | S20 | I | 6 |
| L15 | II | 0 | | S5 | II | 0 |
| L17 | II | 2 | | S7 | II | 1 |
| L23 | II | 7 | | S11 | II | 0 |
| L5 | III | 0 | | S12 | II | 1 |
| L18 | III | 3 | | S9 | III | 1 |
| L29 | III | 21 | | S10 | III | 0 |
| L34 | III | 21 | | S13 | III | 0 |
| L2 | IV | 0 | | S14 | III | 0 |
| L14 | IV | 0 | | S19 | III | 0 |
| L19 | IV | 5 | | S2 | IV | 2 |
| L32 | IV | 16 | | S3 | IV | 0 |
| L6 | V | 0 | | S21 | IV | 197 |
| L9 | V | 15 | | | | |
| L11 | V | 1 | | | | |
| L28 | V | 5 | | | | |
| L33 | V | 7 | | | | |
| L7/L12 | VI | 2 | | | | |
| L10 | VI | 5 | | | | |
| L16 | VI | 0 | | | | |
| L25 | VI | 109 | | | | |
| L27 | VI | 2 | | | | |
| L30 | VI | 78 | | | | |
| L31 | VI | 6 | | | | |
| L35 | VI | 7 | | | | |
| L36 | VI | 2 | | | | |

[a] – Ribosome assembly order and cell coloring is according to Chen and Williamson, 2013 (67)

[b] – Total number of genomes in COGs that are lacking the genes for respective proteins. L29 is mostly lost in tiny genomes (Table 1), absence of L25, L34, and S21 is mostly due to lineage-specific gene loss (Table 2).
